# Simulations of dynamically cross-linked actin networks: morphology, rheology, and hydrodynamic interactions

**DOI:** 10.1101/2021.07.07.451453

**Authors:** Ondrej Maxian, Raúl P. Peláez, Alex Mogilner, Aleksandar Donev

**Affiliations:** Courant Institute, NYU, New York, NY 10012; Department of Theoretical Condensed Matter Physics, Universidad Autónoma de Madrid, 28049, Madrid, Spain; Department of Biology, NYU, New York, NY 10012

## Abstract

Cross-linked actin networks are the primary component of the cell cytoskeleton and have been the subject of numerous experimental and modeling studies. While these studies have demonstrated that the networks are viscoelastic materials, evolving from elastic solids on short timescales to viscous fluids on long ones, questions remain about the duration of each asymptotic regime, the role of the surrounding fluid, and the behavior of the networks on intermediate timescales. Here we perform detailed simulations of passively cross-linked non-Brownian actin networks to quantify the principal timescales involved in the elastoviscous behavior, study the role of nonlocal hydrodynamic interactions, and parameterize continuum models from discrete stochastic simulations. To do this, we extend our recent computational framework for semiflexible filament suspensions, which is based on nonlocal slender body theory, to actin networks with dynamic cross linkers and finite filament lifetime. We introduce a model where the cross linkers are elastic springs with sticky ends stochastically binding to and unbinding from the elastic filaments, which randomly turn over at a characteristic rate. We show that, depending on the parameters, the network evolves to a steady state morphology that is either an isotropic actin mesh or a mesh with embedded actin bundles. For different degrees of bundling, we numerically apply small-amplitude oscillatory shear deformation to extract three timescales from networks of hundreds of filaments and cross linkers. We analyze the dependence of these timescales, which range from the order of hundredths of a second to the actin turnover time of several seconds, on the dynamic nature of the links, solvent viscosity, and filament bending stiffness. We show that the network is mostly elastic on the short time scale, with the elasticity coming mainly from the cross links, and viscous on the long time scale, with the effective viscosity originating primarily from stretching and breaking of the cross links. We show that the influence of nonlocal hydrodynamic interactions depends on the network morphology: for homogeneous meshworks, nonlocal hydrodynamics gives only a small correction to the viscous behavior, but for bundled networks it both hinders the formation of bundles and significantly lowers the resistance to shear once bundles are formed. We use our results to construct three-timescale generalized Maxwell models of the networks.

## 1 Introduction

In most cells, as much as 10% of all protein is actin [62]. The majority of actin is the F-actin cytoskeleton – a gel made of rapidly-turning-over (assembling and disassembling) actin filaments (which we will call fibers), which are dynamically cross linked by a vast host of actin binding proteins (which we will call cross linkers or CLs). The actin cytoskeleton is largely responsible for the cell’s shape, movements, division, and mechanical response to its external environment [2]. Thus, understanding the mechanical and rheological properties of dynamically cross-linked cytoskeletal networks is the first step to understanding the mechanical properties of the cell at large.

Several in vitro experimental techniques, including active poking, parallel plate shearing, and embedded-microbead tracking, have characterized the mechano-rheological behavior of cross-linked actin gels [62]. Depending on the experimental conditions, these studies report viscoelastic moduli in a wide range, from 0.1 to hundreds of Pascals [31, 34, 86, 47]. The mixed viscoelastic mechanical response of the densely cross-linked gel is expected: simply speaking, at short time scales the network of elastic fibers and CLs can be considered permanently interconnected and deform elastically, while at long time scales dynamic CLs are expected to connect fibers only transiently, enabling the brief storage of elastic energy of deformations. This elastic energy is dissipated after the CLs detach, which causes effective viscous behavior on long timescales. Often, the measured elastic modulus for actin gels is about an order of magnitude greater than the viscous one [34, 31, 47]; however, some studies measure elastic and viscous moduli of similar magnitude [11], and in some cases the elastic modulus is smaller than the viscous one [87]. Both moduli are increasing functions of actin and CL concentrations [22, 23]; see especially [23] for master curves over a wide range of CL and actin concentrations.

To understand the intrinsic timescales in the gel, many experimental studies examine the dependence of the loss and storage moduli on the *frequency* of oscillatory shear deformation *ω*. All experimental studies agree that the behavior of transiently cross-linked networks on long timescales (low frequencies) is qualitatively viscous both in vitro [67, 74] and in vivo [11], with many studies reporting a decay rate of *ω*^1/2^ for both moduli at low frequencies in the absence of actin turnover [67, 95, 65]. As the frequency increases, some studies report a monotonically-increasing viscous modulus [23], while others display a curious local maximum and minimum in the viscous modulus data [48, 95, 96]. Proceeding to the high-frequency limit, there are again contradictions, as some studies report a viscous modulus that scales as *ω*^0.75^ [23] (consistent with the thermal fluctuations of a single semiflexible filament [64, 47]), while others observe a viscous-fluid-like scaling of *ω*^1^ [17, 18], and still others yield a scaling of *ω*^1/2^ [67, 74, 100], which is characteristic of the Rouse model of polymer physics (fluctuating beads joined by harmonic springs) [73]. Thus the short and intermediate timescale behavior, and its dependence on microscopic parameters, is still an open question, as is the exact meaning of “long” and “short” timescales (i.e., when these observed scalings begin to dominate).

Part of the reason for the variations in experimental data is that each actin binding protein produces a unique change in network morphology [21], which in turn uniquely affects the viscoelastic moduli. For compliant CLs such as filamin, Kasza et al. observe a strong frequency-dependence of *ω*^0.7^ for the viscous modulus and a weak power law scaling of the elastic modulus, with both taking values in the range 0.1 – 1 Pa [38]. As the CLs get shorter and stiffer, the frequency-dependence in the elastic [38, 87, 23, 88] and sometimes even viscous modulus disappears [38], which suggests a change in the network morphology with the type of cross-linker. It is shown in [67] that changes in the moduli with chemical cross linking are only relevant when the cross-linker length is shorter than the mesh size, because these kind of CLs group the fibers into bundles, thereby changing the macroscopic structure of the network.

Wachsstock et al. [86] study the relationship between mechanical response and morphology using actin filaments with *α*-actinin cross linkers. Their experiments and modeling show a transition from elastic, solid-like behavior to viscous, fluid-like behavior as the *α*-actinin and actin concentrations increase, which corresponds to the formation of bundles within the network. These bundles are no longer entangled in a complex network and are free to slide past each other, which leads to viscous behavior. This observation is compatible with the work of Tseng et al. [84], which shows that more homogeneous networks are more elastic in nature, and suggests that the cell makes the strongest elastic structures by combining bundling and cross-linking proteins to form a cross-linked network of bundles [85].

In view of this experimental complexity, many modeling studies address scaling of the cross-linked actin gel’s mechanical moduli with kinetic and microscopic mechanical parameters. Some of the rheological behavior of pure actin networks can be explained with semiflexible polymer theory [54, 33]. For instance, strain-stiffening behavior in filament networks can be accounted for by an entropic model, in which each thermally fluctuating polymer resists affine (stretching) deformations, or an enthalpic model, in which the filaments bend before they stretch, allowing them to reorganize along the direction of shear [69]. Semiflexible filament theories of this nature have recently been extended to transiently cross-linked networks [10, 96, 66], and it has been shown that a broad spectrum of relaxation times appears for a cross linker with a single unbinding rate, because larger and larger wavelength bending modes are able to relax as cross linkers unbind [10, 65]. These theories may explain the soft glassy rheology observed in living cells [65] and the low-frequency regime where the moduli scale with the square root of frequency [96, 10].

Speaking more broadly, the existing models of cross-linked networks fall into two categories: continuum and agent-based (discrete) models. There are a number of continuum models available for the cell cytoskeleton, and in fact there has been a lot of work in recent years on extending the immersed boundary method [71] to cytoskeleton-like fluids. For example, Karcher et al. employ a continuum finite element model to measure the stress induced by magnetic-bead-forcing [37] (a similar model was also used in [61]). They model the cytoskeleton as either a Maxwell (spring and dashpot in series) or Kelvin-Voigt (spring and dashpot in parallel) material and report a large sensitivity of the mechanical behavior on the choice of model. Other studies have approximated the cell cytoskeleton as a poroelastic [80], porous viscoelastic [12], or Brinkmann [28] fluid. In either case, as discussed in [37, 62], continuum models are only as good as their fits to the data, and they in general have difficulty relating macroscopic parameters (such as porosities, discrete timescales, and stiffnesses) to microscopic parameters. In fact, it has been suggested that the cell possesses a continuum of relaxation timescales [14, 65], which would invalidate any continuum model with a discrete number of elements.

Discrete models tend to suffer the opposite problem in that they are detailed and realistic, but are harder to analyze and do not readily extrapolate to macroscopic systems. Most of these models have focused on simulating the steady state structure of actin networks, which presumably could help explain some of the observed rheological behavior. Head et al. [29], for instance, solve an energy minimization problem for straight filaments in two dimensions and use their results to characterize the transition from affine (stretching-dominated) to non-affine (bending-dominated) deformation in cross-linked networks. In a more detailed approach, Kim et al. [41] use Brownian dynamics simulations on a bead-spring model of actin to show that bundling is reduced when CLs are forced to bind in the perpendicular direction (as they do when fascin is the cross-linker). They propose that the ability of a network to flow on long timescales is due to the breaking of CLs with shear, followed by their reformation on a timescale determined by network reorganization and not individual CL binding. There have also been a number of computational studies on the structure and contractile behavior of *actomyosin* networks, which show that a critical concentration of *α*-actinin cross-linkers can combine with myosin motors to form ordered bundles of actin filaments with varying polarity [72, 46].

Two recent studies [66, 90] combine computational rheology with microscopic Brownian dynamics simulations to measure the rheological properties of transiently cross-linked actin networks *without* filament turnover and without hydrodynamic interactions between the filaments. In these studies, a local maximum in the viscous modulus is observed [48], which was shown to correspond to a maximum in CL binding and unbinding events [90]. However, the networks considered in these studies are either homogeneous [90, Fig. 2] or highly bundled [66, Fig. 1], and we know of no detailed simulations that have reported the rheological properties over a range of structures and microscopic parameters. Furthermore, due to the absence of turnover, in both of these studies the structure being simulated is not in equilibrium and evolves significantly with the imposed flow, and so the measured low-frequency viscous and elastic moduli depend on history and how the measurement is performed, and are consequently not well-defined.

**Figure 1:**
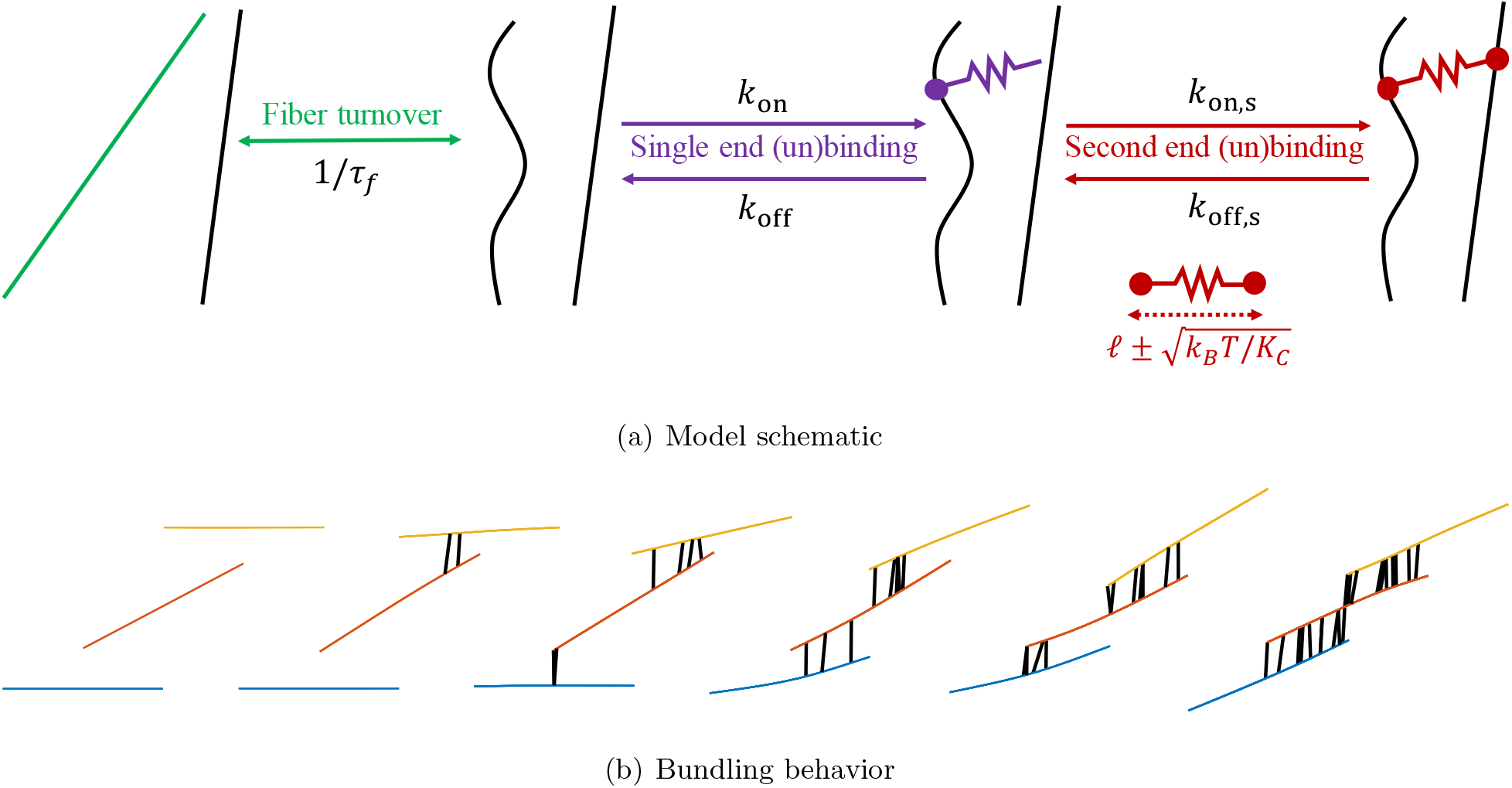
(a) Our model of dynamic cross-linking with fiber turnover. We coarse-grain the dynamics of individual CLs into a rate for each end to bind to a fiber. The first end (purple) can bind to one fiber at any binding site. Once bound, we account for thermal fluctuations of the CL length by allowing the second end (red) to bind to any other fiber within a distance 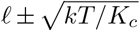 of the first end, where *ℓ* is the CL rest length and *K_c_* is the CL stiffness. To model actin turnover, we allow each fiber to disassemble with rate 1/*τ_f_* and for a new (straight) fiber to assemble in a random place (nascent fiber is shown in green) at the same time. (b) Consecutive simulation snapshots illustrate how the model reproduces the bundling behavior characteristic of an actin mesh cross-linked with *α*-actinin. A pair of fibers that are close enough to be crosslinked by a thermally stretched CL are pulled together when this CL relaxes to its rest length. This brings the fibers closer together, promoting binding of additional CLs. When multiple CLs along the inter-fiber overlap relax to their rest length, cross-linked pairs of fibers align and stay close together, making a bundle.

**Figure 2:**
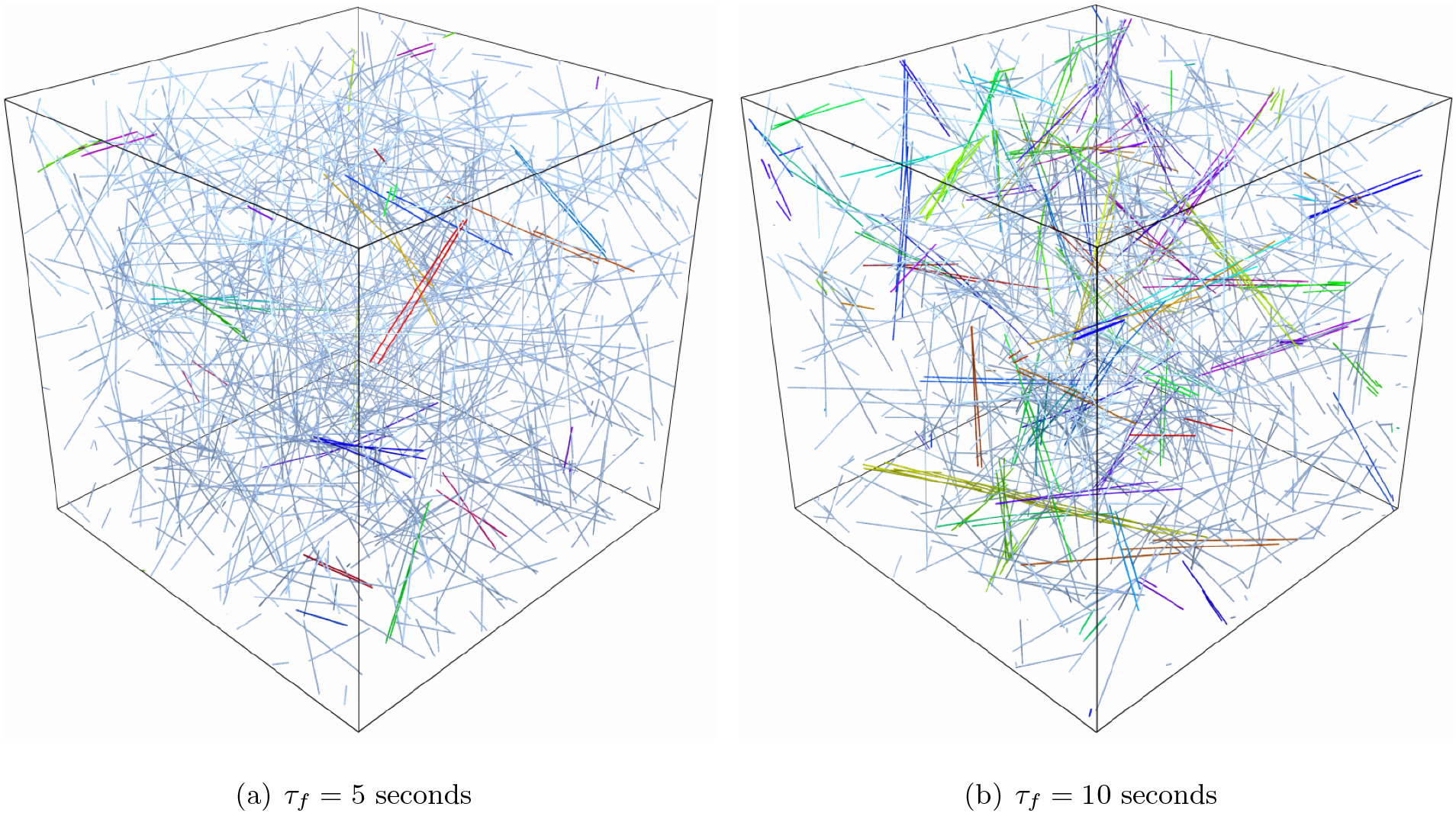
The actin gel for (a) *τ_f_* = 5 seconds and (b) *τ_f_* = 10 seconds, both shown on a domain with edge length *L_d_* =3 *μ*m. Fibers in the same bundle are colored with the same color. In (a), we observe a more homogeneous mesh. In this case, we use *L_d_* = 2 *μ*m in most simulations. In (b), we observe multiple bundles embedded into a mesh, and we use *L_d_* = 3 *μ*m in most simulations.

Despite, and maybe even because of, the large volume of experimental and theoretical studies on cross-linked actin gel mechanics, a number of questions still remain open. We focus on three of them: What role does the morphology of the network play in the mechanical response? Could there be a significant contribution from nonlocal hydrodynamic interactions? And can we inform a simple and practical spring-and-dashpot model by detailed microscopic simulations?

This paper is built around answering these questions. To do so, we first review our formulation for inextensible, semiflexible filaments [57] and extend it to the case of transient cross-linking. We model filament polymerization dynamics by turning over filaments with a characteristic time *τ_f_* and introduce an operator splitting scheme to update the system by turning over the filaments, updating the cross-linked network, and moving the filaments in a sequential order. Since previous experimental and modeling studies show that the mechanical behavior of actin networks is most affected by the presence of short, stiff CLs which induce morphological changes in the network, we will focus our analysis on CLs with spring stiffness 10 pN/*μ*m and a 50 nm rest length, thereby mimicking *α*-actinin. We show that changing the turnover time *τ_f_* with all other parameters fixed gives two model systems: an isotropically cross-linked network, or homogeneous meshwork, and a composite network of bundles embedded in the meshwork (here we follow the classification scheme of [49]). Unlike previous simulation studies [66, 90], the networks that we consider here are in a dynamic steady state, which allows us to precisely quantify their steady-state mechanical properties. In particular, we use small amplitude oscillatory shear (SAOS) rheology over a range of biologically-relevant frequencies (0.01 Hz to 100 Hz)^1^ to show that the networks have three characteristic timescales on which the links go from rigid to flexible, permanent to dynamic, and viscoelastic to purely viscous. It is on this last timescale that remodeling of the network occurs.

This paper is, to our knowledge, the first to incorporate many-body hydrodynamic interactions (HIs) in the study of transiently cross-linked actin networks. Here we once again find results that depend on the timescale under consideration. At high frequency, we show that the viscous modulus scales linearly with *ω*, and that accounting for nonlocal hydrodynamics simply changes this coefficient by a fixed percentage. In contrast, at low frequency, we show that nonlocal hydrodynamic interactions lead to a significant *decrease* in the moduli for structures with a significant degree of bundling, and that the decrease can be explained by the disturbance flows inside a bundle. We conclude this paper by using our knowledge of the timescales and role of nonlocal hydrodynamic interactions to suggest coarse-grained continuum models for the transiently cross-linked actin gel.

## 2 Materials and methods

We first introduce our simulation method for passively cross-linked actin gels. This begins with a brief review of our recently-developed spectral method for semiflexible, inextensible fibers in the presence of cross linkers [57]. We next introduce our models of dynamic cross-linking and actin turnover, both of which are based on the next reaction stochastic simulation algorithm. We then discuss the three approximations to the mobility (force-velocity) relationship: (non-isotropic) local drag, intra-fiber hydrodynamics, and full nonlocal hydrodynamics, as well as how we evaluate them. Most of the details of this can be found in [57]; here we only elaborate on the modifications we have made for efficient simulation of a bundled many-fiber gel. We conclude this section with our temporal integration (network evolution) strategy, which uses operator splitting to turn over the fiber positions, update the cross-linked network, and update the fiber positions in a sequential order.

### 2.1 Semiflexible, inextensible fibers

Since actin fibers are slender and semiflexible, we can represent them by smooth Chebyshev interpolants ***X***(*s*), where *s* ∈ [0, *L*] is an arclength parameterization of the fiber centerline and *L* is the fiber length. We denote the tangent vector by ***τ***(*s*) = ***X**_s_*(*s*). For inextensible filaments such as actin, we have the constraint

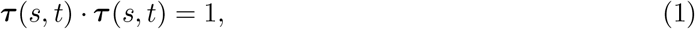

for all *s* and *t*. To enforce the constraint, we introduce a Lagrange multiplier force density **λ**(*s*, *t*) on each fiber.

At every instant in time, each fiber resists bending with bending force density (per unit length)

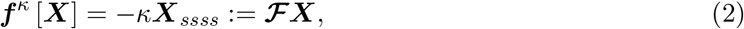

where the constant linear operator 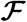 gives ***f**^κ^* with the “free fiber” boundary conditions [83]

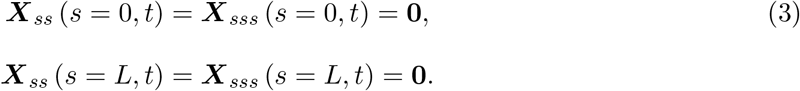

In [57, Section 4.1.3], we discuss how to discretize the operator 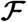 to spectral accuracy in a manner consistent with the boundary conditions (3) using a method called rectangular spectral collocation [6, 16].

Beginning with a single fiber, let us introduce the mobility operator 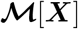 and a spatially and temporally varying background flow ***u***_0_(***x**, t*). Then, accounting for the constraint force, bending force, and any other external forces ***f***^(CL)^ (e.g., those coming from attached CLs), the evolution equation on the fiber is given by

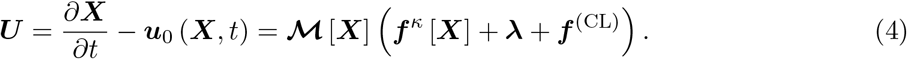

subject to the boundary conditions (3) and inextensibility constraint (1). In [57], we show how to obtain **λ** and enforce inextensibility via the principle of virtual work. We discuss different approximations to 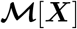 in Section 2.4.

### 2.2 Dynamic cross-linking model

We now specify a method to compute the external or cross-linking force density ***f***^(CL)^ in (4). Here we extend [57] to account for *transient* cross linking, where the links can come on and off through the course of a simulation. To model this, we assume that the density of cross-links is sufficiently high that CLs do not get locally depleted by binding, and that the links diffuse rapidly. By “rapid” diffusion, we mean that the Brownian motion of the links is fast relative to the residence time of a single link, which is on the order of one second (see Table 1). To justify this, we consider the translational diffusion of a rod of length ℓ = 50 nm and radius *a* = 4 nm. The translational diffusion coefficient of the rigid rod can be approximated by Broersma’s relations as [98]

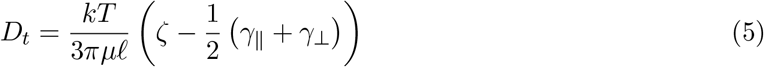

**Table 1.**
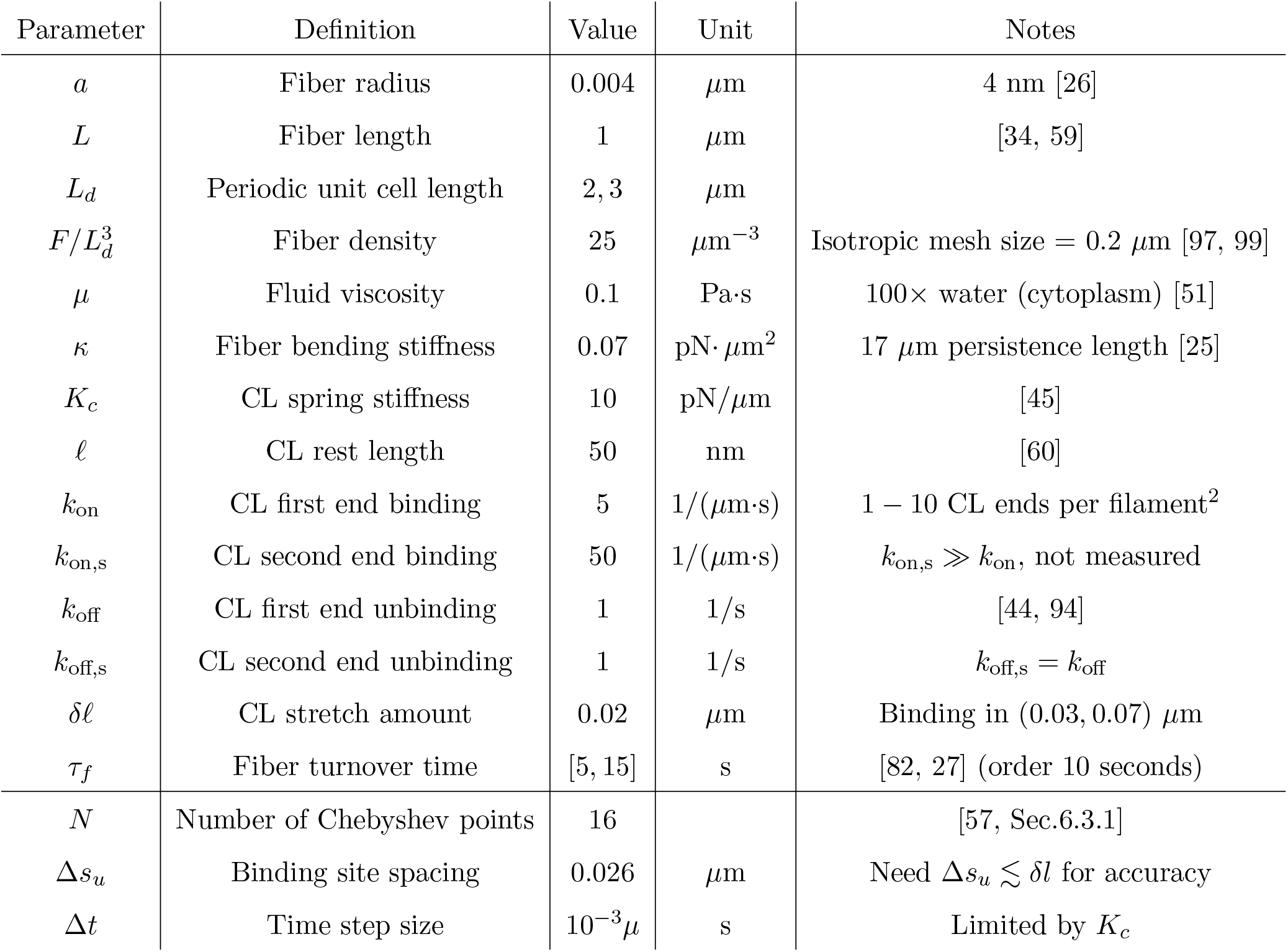
Simulation parameters. With the exception of the changes described in Table 2, we keep these parameters constant in all simulations.

| Parameter | Definition | Value | Unit | Notes |
| --- | --- | --- | --- | --- |
| $a$ | Fiber radius | 0.004 | $\mu\text{m}$ | 4 nm [26] |
| $L$ | Fiber length | 1 | $\mu\text{m}$ | [34, 59] |
| $L_d$ | Periodic unit cell length | 2, 3 | $\mu\text{m}$ | |
| $F/L_d^3$ | Fiber density | 25 | $\mu\text{m}^{-3}$ | Isotropic mesh size = 0.2 $\mu\text{m}$ [97, 99] |
| $\mu$ | Fluid viscosity | 0.1 | Pa·s | 100× water (cytoplasm) [51] |
| $\kappa$ | Fiber bending stiffness | 0.07 | pN·$\mu\text{m}^2$ | 17 $\mu\text{m}$ persistence length [25] |
| $K_c$ | CL spring stiffness | 10 | pN/$\mu\text{m}$ | [45] |
| $\ell$ | CL rest length | 50 | nm | [60] |
| $k_{\text{on}}$ | CL first end binding | 5 | 1/($\mu\text{m}\cdot\text{s}$) | 1 – 10 CL ends per filament<sup>2</sup> |
| $k_{\text{on},s}$ | CL second end binding | 50 | 1/($\mu\text{m}\cdot\text{s}$) | $k_{\text{on},s} \gg k_{\text{on}}$, not measured |
| $k_{\text{off}}$ | CL first end unbinding | 1 | 1/s | [44, 94] |
| $k_{\text{off},s}$ | CL second end unbinding | 1 | 1/s | $k_{\text{off},s} = k_{\text{off}}$ |
| $\delta\ell$ | CL stretch amount | 0.02 | $\mu\text{m}$ | Binding in (0.03, 0.07) $\mu\text{m}$ |
| $\tau_f$ | Fiber turnover time | [5, 15] | s | [82, 27] (order 10 seconds) |
| $N$ | Number of Chebyshev points | 16 | | [57, Sec.6.3.1] |
| $\Delta s_u$ | Binding site spacing | 0.026 | $\mu\text{m}$ | Need $\Delta s_u \lesssim \delta\ell$ for accuracy |
| $\Delta t$ | Time step size | $10^{-3}\mu$ | s | Limited by $K_c$ |

For our parameters in Table 1, we obtain *ζ* = 2.53, *γ*_‖_ = 1.25, *γ*_⊥_ = 0.19, and *D_t_* = 0.15 *μ*m^2^/s. This implies a center of mass displacement of 〈*r*^2^〉 = 6*D_t_t* ≈ 0.9*t*. Thus the time for a CL to travel a mesh size of *r* = 0.2 *μ*m is about 0.04 seconds, which is much faster than the one second lifetime of each link. These assumptions considerably simplify our simulations, as we do *not* have to keep track of explicit CLs, but only the number of links bound to each fiber, which can be tuned by setting the ratio of the binding and unbinding rates.

There are four reactions in the model (parameters are summarized in Table 1):

1. The attachment of one end of a free CL to a fiber, which occurs with rate *k*_on_ (units 1/(length×time)). Here *k*_on_ accounts for the rate of diffusion of the CL until one end gets sufficiently close to a fiber, and then the reaction for the CL to bind to the site.
2. The detachment of a singly-bound CL end to become a free-floating CL, which occurs with rate *k*_off_ (1/time). This is the reverse of reaction 1.
3. The attachment of the second end of a singly-bound CL to *another* (distinct) fiber, which occurs with rate *k*_on,s_ (units 1/(length×time)). This necessarily follows after reaction 1. The rate *k*_on,s_ accounts for the diffusion of the second end to find a fiber within a sufficient distance, and then the binding reaction for the end to latch on to the fiber. We model the thermal fluctuations in the CL length by allowing a singly-bound CL to bind its free end to another attachment site on a different fiber that is within a distance interval

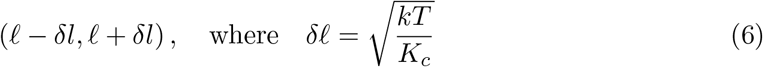

and *kT* ≈ 4 × 10^−3^ pN · *μ*m. In this paper, we will fix *K_c_* = 10 pN/*μ*m [45], so that the distance a link can stretch is *δ*ℓ = 0.02 *μ*m. This is 40% of the rest length of ℓ = 0.05 *μ*m.
4. The detachment of a single end of the doubly-bound CL, so that it becomes singly-bound, which occurs with rate *k*_off,s_ (1/time). This is the reverse of reaction 2.

A schematic of the four reactions is shown in Fig. 1(a). In our model, we do not account for any dependence of the binding/unbinding rates on the CL stretch.

To simulate these reactions stochastically, we employ a variant of the Stochastic simulation/ Gillespie algorithm [24, 4]. To advance the system by a time step Δ*t*, we calculate the rate of each reaction and use the rate to sample an exponentially-distributed time for each reaction to occur. We choose the minimum of these times, advance the system to that time, and then recalculate the rates for the reactions that were affected by the previous step. We organize the event queue efficiently using a heap data structure [15], so that we can compute the next reaction in 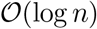 time.

We break each filament into *N_s_* possible uniformly-spaced binding sites, where each of the sites represents a binding region of size *L*/(*N_s_* – 1) = Δ*s_u_*. To properly resolve the possible connections between fibers, we require Δ*s_u_* ≲ *δ*ℓ. We then compute the rates of each reaction in the following way:

1. Single end binding: the rate of a single end binding to one of the *N_s_* attachment points on one of the *F* fibers is *k*_on_Δ*s_u_*. We implement this as a single event with rate *N_s_Fk*_on_Δ*s_u_*, where the specific site is chosen uniformly at random once the event is selected. Multiple CLs are allowed to bind to a site, since each site represents a fiber segment of length Δ*s_u_* that is typically longer than the minimal possible distance between two CLs.
2. Single end unbinding: we keep track of the number of bound ends on each of the *N_s_F* sites. If a site has *b* ends bound to it, we schedule a single end unbinding event from that site with rate *k*_off_*b*. We do this for each site separately, so that this reaction contributes *N_s_F* events to the event queue.
3. Second end binding: we first construct a neighbor list of all possible pairs of sites (*i, j*) that are on distinct fibers and separated by a distance ℓ + *δ*ℓ or less, using a linked list cell data structure [3]. If site *i* has *b* free ends bound to it, we schedule a binding event for that pair of sites with rate *bk*_on,s_Δ*s_u_*. Notice that this naturally implies a zero binding rate if there are no free ends bound to site *i*. We treat the (*j, i*) connection separate from the (*i, j*) one for simplicity at the cost of increasing the number of events in the queue.
4. Second end unbinding: if a link exists between sites *i* and *j*, it can unbind from one of the sites with rate *k*_off,s_. Letting *β* be the number of links in the system, the total rate for which bound links unbind from one end is 2*βk*_off,s_. We therefore schedule a single event with rate 2βk_off,s_, and, once it is chosen, select the link and end to unbind randomly and uniformly.

In the rest of this paper, we will denote the connectivity of the network (number of single ends bound to each site, list of sites connected by CLs) as ***Y***(*t*). The CL force in (4) is then a function of the fiber positions and network configuration, which we denote as ***f***^(CL)^ (***X***; ***Y***).

We use our previous work [57, Section 6.1] to define the cross-linking force density between two fibers ***X***^(*i*)^ and ***X***^(*j*)^ in a manner that can preserve the problem smoothness and therefore the spectral accuracy of our numerical method. Suppose that a CL connects two fibers by attaching to arclength coordinate 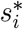 on fiber *i* and 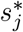 on fiber *j*. Then the force density on fiber *i* due to the CL is

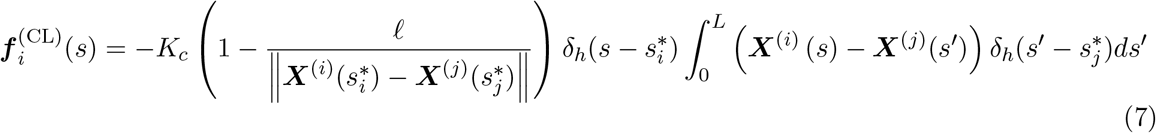

where *K_c_* is the spring constant for the CL (units force/length), ℓ is the rest length, and *δ_h_* is a Gaussian density with standard deviation *σ*. As discussed in [57, Section 6.1], the Gaussian width *σ* depends on how many points are used to discretize the fibers. In this paper, we use *N* = 16 points in the fiber discretization, and so we also use the required *σ*/*L* = 0.1 to preserve smoothness. In the limit *σ* → 0, the Gaussian density becomes a Dirac delta function, and the force density (7) becomes the expected expression for a linear spring connecting the points 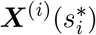 and 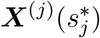. This model is of course an approximation; as shown in [45], the stress-strain relationship for *α*-actinin, which is the model CL we consider here, is nonlinear and exhibits hysteresis for loading and unloading. We follow a number of other modeling studies [90, 42] in approximating *α*-actinin as a linear spring, although we do not include an angular stiffness, since recent experimental results [13] indicate that the dynamics of the *α*-actinin-actin bond are insensitive to rotation.

### 2.3 Fiber turnover

We account for the turnover of actin filaments by removing a filament (and all of its attached links) from the system and rebirthing a new, straight filament with a random location and orientation elsewhere in the system (see schematic in Fig. 1(a)). Our reasoning for this is two-fold: first, it is simple to implement, and second, our simplifying assumptions are supported by experimental data. Experiments have shown that pointed-end depolymerization from one end of an actin filament is too slow to explain the observed rates of filament turnover in vivo [36], and that depolymerization actually occurs in bursts as filaments abruptly sever into smaller pieces [43]. In this sense, our model of depolymerization as an instantaneous event is supported by the data. For polymerization, the process is normally linear, with monomers being added at the plus end until it is capped. It has been shown experimentally [56], however, that the combined time of growth, capping, and disassembly is still significantly smaller than the time in which the filament is at a finite length, so we assume the former to be instantaneous relative to the latter. We refer to [68, Sec. 2.5] for a mathematical treatment of linear polymerization.

Having chosen this model, fiber turnover can be added as another reaction in our stochastic simulation algorithm. Defining the mean turnover time of each fiber as *τ_f_*, we add a fiber turnover reaction to our reaction list with rate *F/τ_f_*. If this reaction is chosen, we select the fiber to turnover randomly and uniformly. We emphasize that our time steps are at most 10^−3^ seconds while the turnover times are of the order seconds, so we almost always see zero or one fiber turnover events per time step.

### 2.4 Mobility evaluation

We complete our description of the evolution equation (4) by specifying the mobility operator 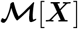. Our approach is fully described in [57, Sec. 2, Appendix A] and is based on an improved version of the traditional slender body theory for Stokes flow [39, 35, 83]. For a single fiber, the self-mobility gives the fiber velocity in (4) as

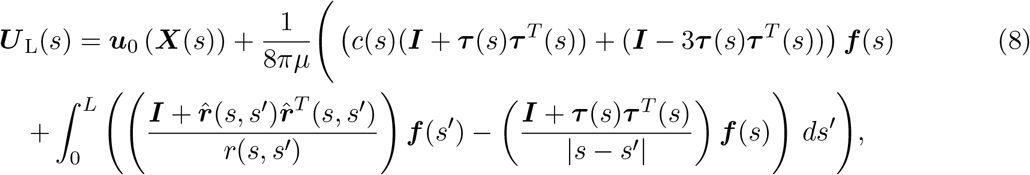

where ***r***(*s*, *s*′) = ***X***(*s*) – ***X***(*s*′), *r* = ‖*r*‖, and 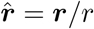. Thus the velocity on the fiber can be written as a leading order local drag term (the first line), which accounts for forcing localized around ***X***(*s*), plus a “finite part” integral remainder term (the second line), which accounts for the intra-fiber *nonlocal* hydrodynamic interactions. In (8), *c*(*s*) is a local drag coefficient which has a logarithmic dependence on the fiber radius *a* as

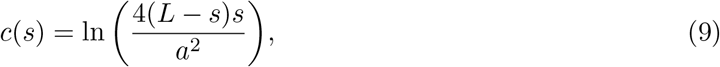

away from the fiber endpoints. Near the endpoints, (9) as written is singular, and we regularize it over a distance *δ* = 0.1*L* as discussed in [57, Section 2.1]. Throughout this paper, we will fix *a* = 4 nm [26].

When there are multiple fibers, the velocity (8) has to be modified to account for the flows induced by fibers *j* ≠ *i* on fiber *i*. Denoting this flow by ***U***_NL_ and indexing the ith fiber with a superscript (*i*), we use a line integral of the Rotne-Prager-Yamakawa (RPY) hydrodynamic tensor over all other fibers to obtain the total disturbance flow

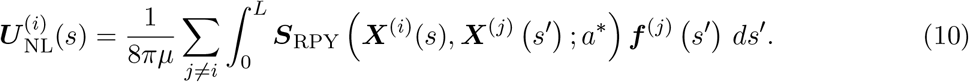

The RPY tensor ***S***_RPY_, which involves the Stokeslet and doublet singularities of Stokes flow, is a specific form of a symmetrically regularized Stokeslet, as explained in more detail in [57, Sec. 2] and [89]. The constant *a*\* is proportional to the fiber radius *a*, and is chosen to match the slender body theory (8) with the RPY integral for a single fiber [57, Eq. 15] (note that in that equation *a* = *ϵL*).

Our strategy for evaluating the total velocity 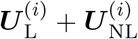 on each fiber is documented in [57, Section 4]. The most computationally intensive part of this calculation is the evaluation of the nonlocal integrals (10). These integrals present challenges because they naively cost 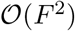 operations to evaluate, where *F* is the number of fibers, and because they can be nearly singular when fibers are close together. For triply periodic systems, which we consider in this paper, we reduce the cost to 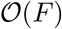 using the positively split Ewald method of [19, 20]. For a summary of how this method is adapted to sheared domains, see [57, Section 4.3].

In our previous work, we dealt with the nearly singular nature of the integrals for nearly contacting fibers by correcting the result from Ewald splitting with the special quadrature method of [1]. While this is the most accurate way to evaluate (10), our recent experiments show that the fastest way to compute the integrals (10) with sufficient accuracy is via oversampled direct quadrature on a GPU. Specifically, we discretize *all* integrals for *j* ≠ *i* using Clenshaw-Curtis quadrature with weights ***w***,

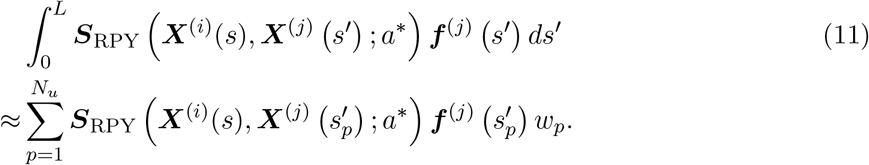

We then perform the summation with periodic boundary conditions using the positively split Ewald method [19, 20, 57], which we implement on a GPU in the UAMMD library [70], with *N_u_* sufficiently large to resolve all of the integrals in (10).

Following [57], to investigate the role of nonlocal hydrodynamic interactions, we vary the model used to compute the fluid velocity on the fibers. Our first option, which we refer to as the *local drag* model, is to include only the first line of (8). Note that this mobility is non-isotropic, since the *c*(*s*) term in (8) gives approximately twice the velocity for a force in the tangential direction, so even the local drag model we use is a significant improvement over formulations based on an isotropic local drag coefficient [78, 90], which are also missing the logarithmic dependence in (9). Another option is to include only the ***U***_L_ term in each fiber’s velocity so that (nonlocal) hydrodynamic interactions between a fiber and itself (which for dilute suspensions are the most important) are accounted for, but not interactions between a fiber and the others. We term this the *intra-fiber* mobility. Finally, when all terms ***U***_L_ + ***U***_NL_ are included on each fiber, we refer to the mobility as the full (nonlocal) hydrodynamic mobility.

### 2.5 SAOS rheology

To quantify the viscoelastic behavior of the network, we measure the linear elastic (also called storage) modulus *G*′ and viscous (loss) modulus *G*″. We employ small-amplitude oscillatory shear (SAOS) rheology to do this [63, 57], so that we impose a shear flow of the form

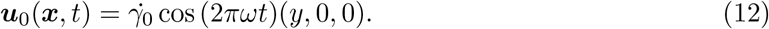

The nonzero component of the *rate* of strain tensor for this flow is given by

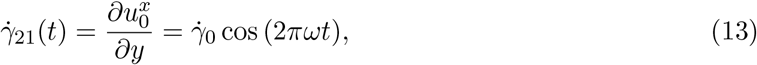

while the nonzero component of the *strain* tensor is

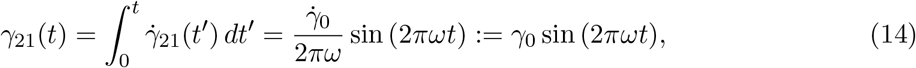

where 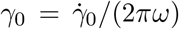 is the maximum strain in the system. The bulk elastic (*G*″) and viscous modulus (*G*″) relate the stress to the strain (for the elastic modulus *G*′) and rate of strain (viscous modulus *G*″) [63] via

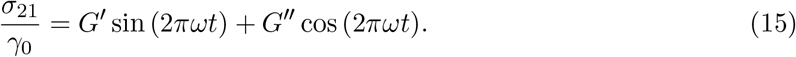

As discussed in [57, Sec. 6.2], the stress tensor for our system can be decomposed into a part coming from the background fluid and a part coming from the internal fiber stresses,

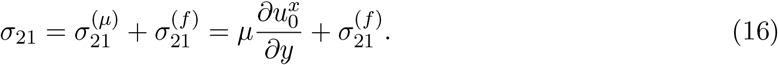

Substituting the background flow (12), we obtain the stress for the background (pure viscous) fluid

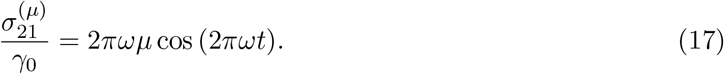

A pure viscous fluid therefore has viscous modulus scaling linearly with *ω* as *G*″(*ω*) = 2*πωμ*. This will be useful to us when we examine the behavior of the viscous modulus of the actin gel.

The contribution of the fibers to the stress tensor is the integral ∫ ***X***(*s*)***f***(*s*) *ds*, where ***f*** is the total force density on the fibers [8, 83], including the elastic forces exerted by the cross-linkers. This must be handled carefully in periodic boundary conditions, as the correct periodic copy of the fiber must be used to ensure the stress is symmetric. The formula we use is given in [57, Eqs. (124,125)].

### 2.6 Temporal integration and splitting

We have three different events to process at each time step: fiber turnover, link turnover, and fiber movement. We will treat them sequentially using a first order Lie splitting scheme [32]. For each time step *n* of duration Δ*t*, from time *n*Δ*t* to (*n* + 1)Δ*t*, we

1. Turnover filaments to update configuration 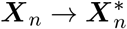.
2. Process binding/unbinding events to evolve 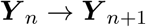.
3. Use the method of [57] to evolve 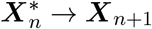.

The evolution of the fiber network in step 3 is performed using the method of [57, Sec. 4.5]. This method obtains second-order accuracy and unconditional stability for fibers with permanent CLs using a combination of a linearly-implicit trapezoidal discretization for the fiber bending force and extrapolations for arguments of nonlinear terms (e.g., we use ***X***_*n*+1/2,*_ = 3/2***X**_n_* – 1/2***X***_*n*–1_ as the argument for the mobility and apply the nonlocal part of the mobility to an extrapolated constraint force **λ**_*n*+1/2,*_ = 2**λ**_*n*–1/2,*_ – **λ**_*n*–3/2_). We use this temporal discretization wherever it applies and is sufficiently stable.

However, extrapolations such as ***X***_*n*+1/2,*_ are nonsensical for fibers that are turned over in the previous time step. For those fibers, we set 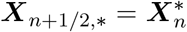 and compute a guess constraint force **λ**_*n*+1/2,*_ by solving (4) with the local drag mobility on each fiber that is respawned. Additionally, in our temporal discretization [57, Eq. (102)], the forcing inside the nonlocal mobility is treated entirely explicitly, since there we assume that the local drag part dominates the fiber motion. There are two ways this could go wrong: first, if a uniform, isotopic suspension is sufficiently concentrated, the temporal integrator becomes unstable. This was the case we already dealt with in [57], where we switched to implicit treatment of the bending force in all parts of the mobility and used an iterative solver. When the fibers are in bundles, however, we find that the approach of [57] is prohibitively expensive due to the ill-conditioning of the elastic force operator 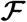 (which involves fourth derivatives) combined with the extrapolations for ***X***_*n*+1/2,*_ and **λ**_*n*+1/2,*_. For this reason, we switch to a first-order backward Euler discretization of fiber elasticity, where ***X***_*n*+1/2,*_ is replaced by 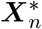 and the elastic force is computed on ***X***_*n*+1_. The backward Euler discretization is our preferred one for simulations with *significant bundling and full hydrodynamics*.

Finally, we discretize the stress tensor in a manner consistent with the numerical method, either as 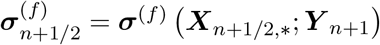 or as 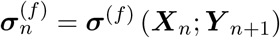. The moduli *G*′ and *G*″ are then evaluated by integrating the stress against the orthogonal sine and cosine functions [57, Eq. (126)].

## 3 Results

This section presents our results for the viscoelasticity of dynamically cross-linked actin networks. We use the parameters in Table 1 to perform the simulations, and show in Section 3.1 that changing the fiber turnover time gives a pair of dynamic steady states with varied degrees of bundling. In Section 3.2, we use a stress relaxation test to show that the networks relax to a state of zero stress on a timescale of a few seconds. This is quantified more precisely with our rheological data in Sections 3.3–3.6, where we establish three discrete relaxation timescales and discuss the behavior on short, long, and intermediate timescales in more detail. Our discussion of the role of nonlocal hydrodynamic interactions (HIs) necessarily follows this, as the role of HIs is more significant on long timescales than short ones, as we show in Section 3.7. We conclude this section by developing a continuum Maxwell-type model that is informed by our knowledge of the three timescales.

Our simulations use a background fluid with viscosity *μ* = 0.1 Pa·s, mimicking the larger viscosity of the cytoplasm [51]. We will, however, assume the background fluid is Newtonian, in contrast to actual cytoplasm, so that we can isolate the contribution of the fibers to the viscoelastic moduli.

### 3.1 Fiber turnover creates dynamic steady states with varied degrees of bundling

We first show that varying the turnover time *τ_f_* of the *F* fibers creates a set of dynamic steady states with varying degrees of bundling. We also verify that the statistics are independent of the system size by considering three possible cubic periodic unit cells of different edge length *L_d_*, all with the same mesh size (computed assuming that the fibers are distributed isotropically) 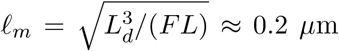. The fibers have length *L* = 1 *μ*m throughout, so we will consider domains of size *L_d_* = 2 *μ*m (*F* = 200 fibers), *L_d_* = 3 *μ*m (*F* = 675 fibers), and *L_d_* = 4 *μ*m (*F* = 1600 fibers). We will show that simulations of a homogeneous meshwork (*τ_f_* = 5 seconds) are suitably carried out in the smallest of the three systems, while runs where bundling is more considerable require the next-largest system (*L_d_* = 3 *μ*m).

As shown in Fig. 1(b), dynamic cross-linking leads to bundling of fibers, a phenomenon that is well documented experimentally for a variety of CL types [50, 76, 22]. To define a bundle, we use the network ***Y*** to map the fibers to a connected graph [52]. Unlike [52], we say that two fibers are connected by an edge if they are connected by two links a distance *L*/4 apart, the idea being that the fibers are sufficiently aligned in that case for the links to constrain their orientations. “Bundles” are then the connected components of this graph (two or more fibers per bundle). To better understand the network morphology, we define a bundle density 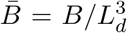 (units bundles/*μ*m^3^), where *B* is the total number of bundles, and also quantify the percentage of fibers in bundles. We define 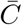 as the average number of links bound to each fiber (the total number of links is 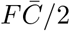).

#### 3.1.1 Homogeneous meshwork morphology

Beginning with turnover time *τ_f_* = 5 s, we perform a steady state run to 10*τ_f_* = 50 s using the three domain sizes. As shown in Fig. 2(a), the network is made primarily of fibers oriented isotropically (as can be verified by computing an order parameter of the type [52, Eq. (21)]), with a few bundles of at most two to three aligned fibers. We quantify this more precisely in Table 2, where we see that there is on average one bundle per *μ*m^3^, and that less than 10% of the fibers are in bundles. Because there are only a small number of bundles and no permanent structures over long timescales in this system, we classify this system as a *homogeneous meshwork*. We will report results for it using a domain size of *L_d_* = 2 *μ*m (we have verified that using *L_d_* = 3 *μ*m does not change the results significantly; see S1 Fig.).

**Table 2.** Description of the systems we consider and their respective computed mean link and bundle densities. These data are from rheology simulations using *ω* = 1. Here 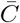 is the average number of links attached to each fiber, 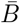 is the bundle density in bundles per *μ*m^3^, and “% in B” is the percentage of the fibers that are in a bundle in the dynamic steady state.

| System | $\tau_f$ (s) | Varied params | $\bar{C}$ | $\bar{B}$ ($\mu\text{m}^{-3}$) | % in B |
| --- | --- | --- | --- | --- | --- |
| Meshwork | 5 | — | 5.8 | 1.0 | 8.5 |
| Meshwork, $\kappa/10$ | 5 | $\kappa = 0.007$ | 5.6 | 0.93 | 8.1 |
| B-In-M | 10 | — | 12.5 | 2.2 | 23 |
| B-In-M, $2k$ | 5.7 | $k_{\text{on}} = 10, k_{\text{on},s} = 100, k_{\text{off}} = k_{\text{off},s} = 2$ | 12.5 | 2.1 | 21 |
| B-In-M, $10\mu$ | 16 | $\mu = 1$ | 12.2 | 1.8 | 17 |

#### 3.1.2 Bundles embedded in meshwork morphology (B-In-M)

In Fig. 2(b), we show snapshots from the dynamic steady state with a (doubled) turnover time of *τ_f_* = 10 s. We observe significantly more bundling and fiber alignment, as well as bundles with several (four to six) fibers in them. In Table 2, we see that the steady-state link density has more than doubled from the 5 s turnover case, and that the percentage of fibers in bundles has gone from about 10% for *τ_f_* = 5 s to 25% for *τ_f_* = 10 s. The fluctuations of link density in the smaller system (*L_d_* = 2) are quite large (about 20%, see S2 Fig.), which makes averaging too inaccurate. For this reason, for *τ_f_* = 10 seconds we run a larger system with *L_d_* = 3 *μ*m, which we show in Fig. 2(b). We use the *L_d_* = 4 system with 1600 fibers to verify that the finite system size has little effect; see, for example, Table 3.

**Table 3.**
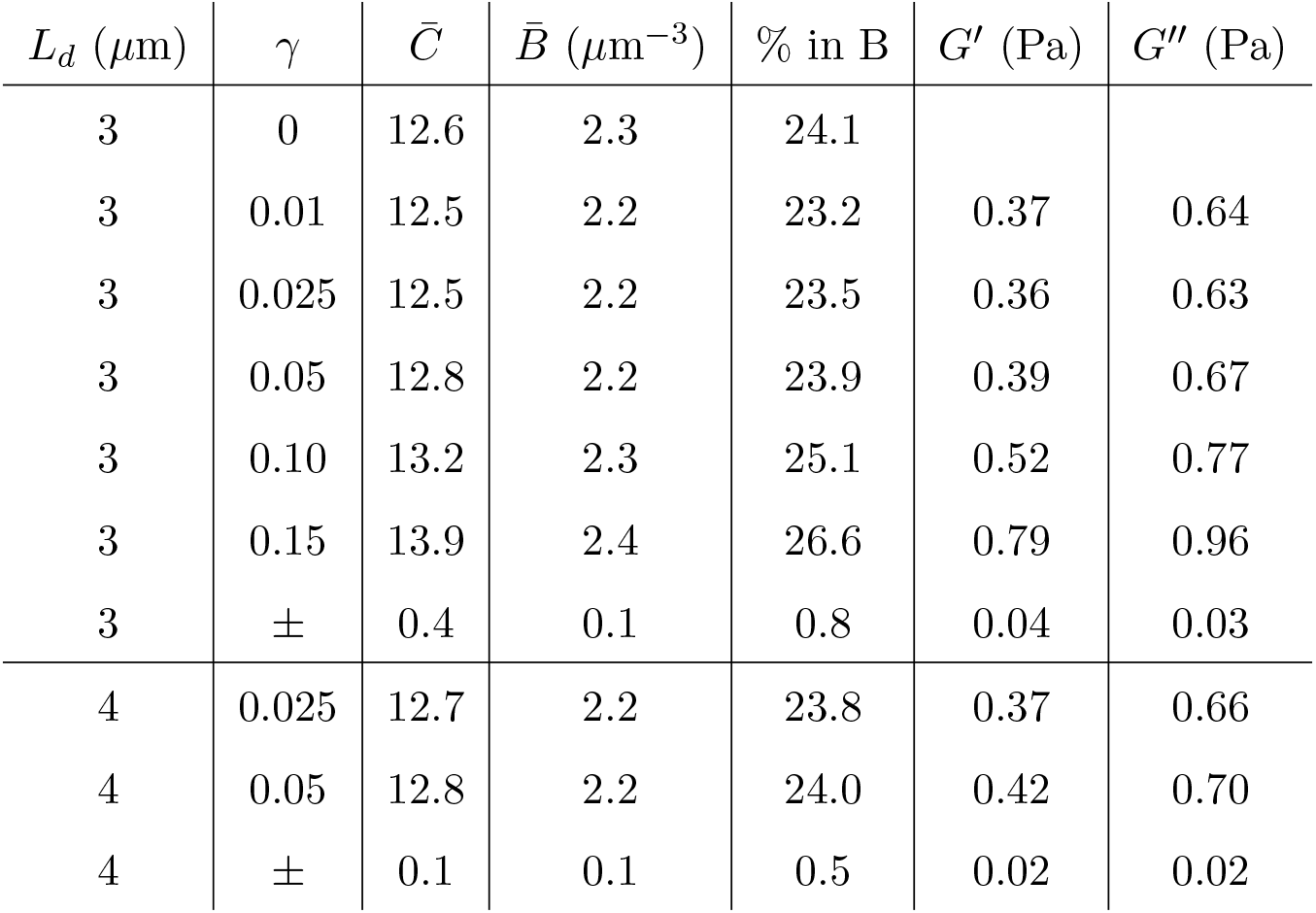
Network structure statistics for the runs over 120 seconds for the B-In-M network with *ω* = 1 Hz. We give the mean for each statistic, with uncertainties on the measurements (± row) equal to two standard deviations across five trials. The uncertainties in the viscoelastic moduli for *γ* = 0.15 are larger, *G*′ = 0.79 ± 0.09 Pa and *G*″ = 0.96 ± 0.12 Pa. The comparable values of the statistics and moduli for *L_d_* = 3 *μ*m and *L_d_* = 4 *μ*m demonstrate that the finite-size errors are smaller than the statistical errors.

Because this system has a significant number, but not more than half, of the fibers in bundles, and because the maximum bundle size is still at most ten fibers, we refer to this system as a *bundles embedded in meshwork (B-In-M*) morphology, where there are small bundles of a few fibers embedded in an otherwise homogeneous mesh.

#### 3.1.3 Other systems (parameter variations)

In addition to varying the morphology through the turnover time, we also look at systems with the same morphologies, but different microscopic parameters. Table 2 gives our description of these systems. For a homogeneous meshwork, we consider *τ_f_* = 5 s again, but this time with ten times smaller bending stiffness, so that *κ* = 0.007 pN·*μ*m^2^. Table 2 shows that the morphology in this case is the same as the homogeneous meshwork with κ = 0.07 pN·*μ*m^2^, so changing the bending stiffness has a minimal impact on the morphology. Indeed, the fibers remain relatively straight even with the smaller bending stiffness (see S3 Fig.), just as they are straight in both the B-In-M and homogeneous meshwork systems in Fig. 2. This implies that the degree of cross-linking/network connectedness is insufficient to induce filament bending, even in B-In-M geometries.

We also consider B-In-M systems with larger viscosity, *μ* = 1 Pa·s instead of 0.1, and another system where we double the rates of link binding and unbinding. Table 2 shows how we vary the turnover time in these systems to obtain a B-In-M morphology. When the viscosity is increased by a factor of 10, we only need to increase the turnover time by a factor of 1.5 to obtain a similar morphology (match 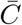 as closely as possible). Meanwhile, doubling the rates forces us to cut the turnover time almost in half to obtain the same morphology. This provides our first indication that the morphology, or the timescale on which the network organizes itself into bundles, is strongly dependent on the link binding and unbinding rates and weakly dependent on the underlying fluid viscosity. This combination of behavior suggests we are operating in a regime where the CLs are attached for long enough to move the fibers into a quasi-steady state (since using twice the link turnover rate speeds up the process by a factor of two and making the dynamics slower by changing viscosity has little effect).

#### 3.1.4 The linear response regime (LRR): viscoelastic moduli and stress spectra

Our goal for the rheology experiments is to measure the shear moduli of the network in its dynamic steady state, and in the linear response regime (LRR). In this section we briefly demonstrate how to find the LRR and what happens when we go beyond it, using the B-In-M geometry as an example.

Table 3 shows the steady state statistics and mean viscoelastic moduli for the B-In-M system with varying strains *γ*. In Table 3 and throughout this paper, we report the viscous modulus *G*″ *without* the contribution from the viscosity of the Newtonian solvent 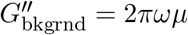. We observe from the table that the LRR is *γ* ≲ 0.05, since using *γ* > 0.05 leads to significantly larger moduli. In particular, Table 3 shows that larger strains of *γ* ≥ 0.10 induce higher bundle densities and the formation of more links. Furthermore, the links that do form in the larger-strain systems are themselves more strained, as the mean square link strain at *γ* = 0.1 and *γ* = 0.15 is 10% and 25% higher, respectively, than in the linear regime. The combination of these two factors gives large increases in the moduli when we reach the nonlinear regime. The viscoelastic moduli for larger strains are also subject to significantly larger uncertainties (e.g., the uncertainity for *γ* = 0.15 is at least double the uncertainity for *γ* = 0.1). The larger uncertainties come from fluctuations in the underlying microsctructure, as the fluctuations in the link density increase by 50% from *γ* = 0.01 to *γ* = 0.15.

Another way we can confirm that we are in the LRR is by looking at the Fourier spectrum of the stress. Specifically, we will write the stress as

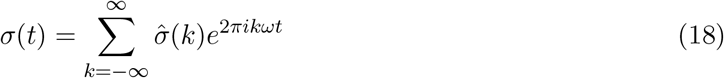

and use the discrete Fourier transform to look at the amplitude of the coefficients 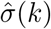 for various *γ*. In Fig. S4, we show the spectrum 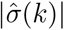 for integer values of *k* with *τ_f_* = 10 s, where we observe a peak at the obvious location of *k* = 1. It is the emergence of higher harmonics for larger strains, however, that tells when we leave the LRR. From S4 Fig., we see that *γ* = 0.1 is definitely *not* in the linear regime for the B-In-M geometry, since it has a *k* = 3 Fourier coefficient that is only about ten times smaller than the one for *k* = 1. Increasing *γ* to 0.15, we see an even stronger response from the *k* = 3 harmonic, and the emergence of the *k* = 2 harmonic as well.

### 3.2 Timescales of stress relaxation

We first seek to gain some understanding of the timescales in the system with a stress relaxation test, snapshots from which are shown in Fig. 3. We simulate for 0.25 seconds using *ω* = 1 Hz, so that the system ends at its maximum strain *γ* (the results are largely independent of both the strain and shear rate used to obtain it). In Fig. 4, we show the decay of the stress, with *t* = 0 denoting the point at which the system reaches the maximum strain. In addition to our dynamic network geometries, for which we average over 30 trials, we consider *permanently* linked, interconnected networks of the type considered in [57], where 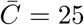 ensures the network is well above the rigidity percolation threshold (over 95% of the fibers are in one connected component, where two fibers are connected if they are linked by at least one CL). These networks, for which we *do not* include fiber turnover, give smoother stress profiles, and so we only average over 5 trials.

**Figure 3:**
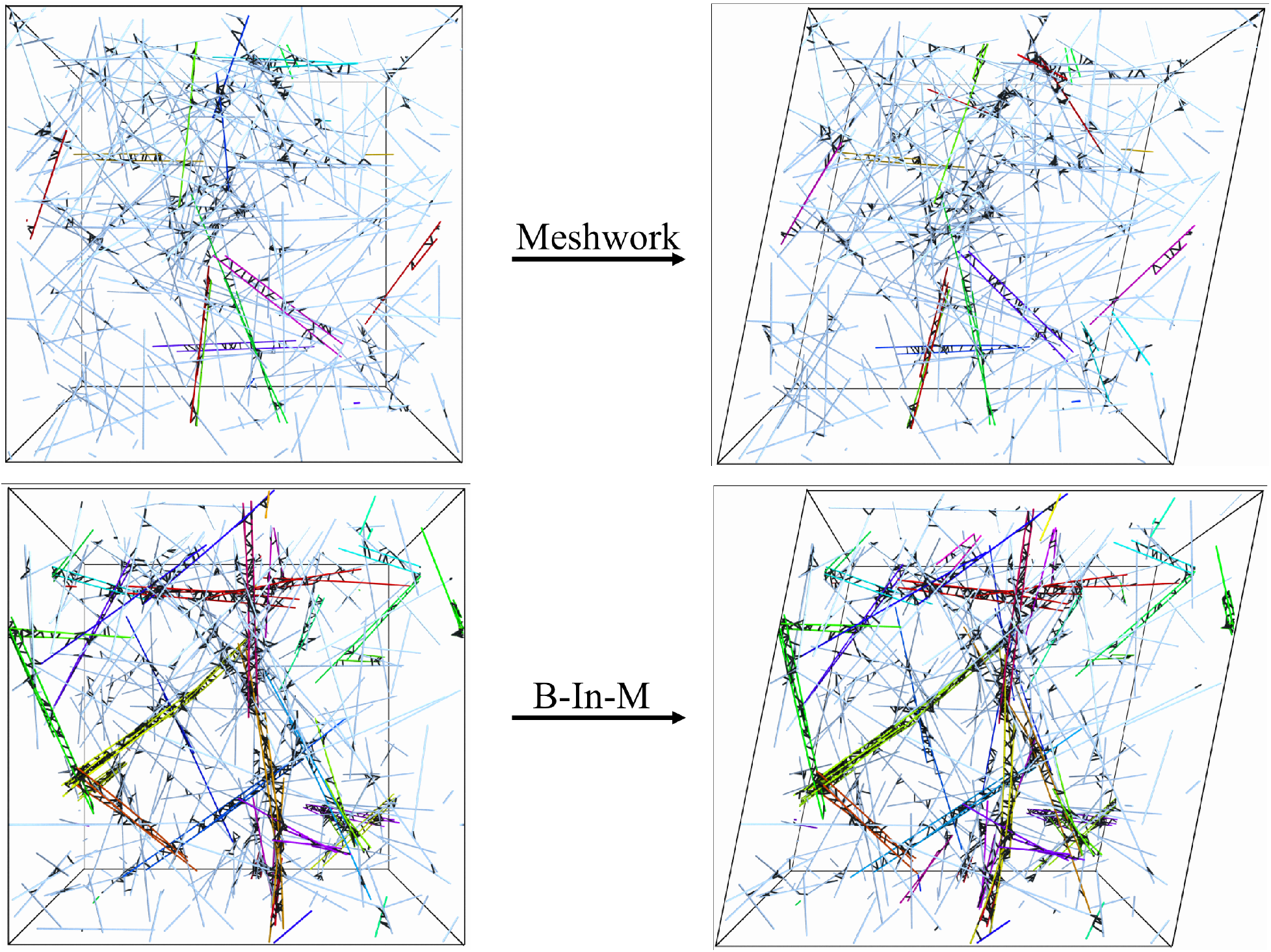
Snapshots from the stress relaxation test in a homogeneous meshwork (top) and B-In-M geometry (bottom). We begin with an unsheared unit cell at left, then shear the network until it reaches a maximum strain (20% in this case, shown at right), after which we turn off the shear and measure the relaxation of the stress. As in Fig. 2, the colored fibers are in bundles, and the CLs are shown in black. For the B-In-M geometry, these snapshots are from a smaller domain (*L_d_* = 2 *μ*m) than we typically use so that we can also see the CLs.

**Figure 4:**
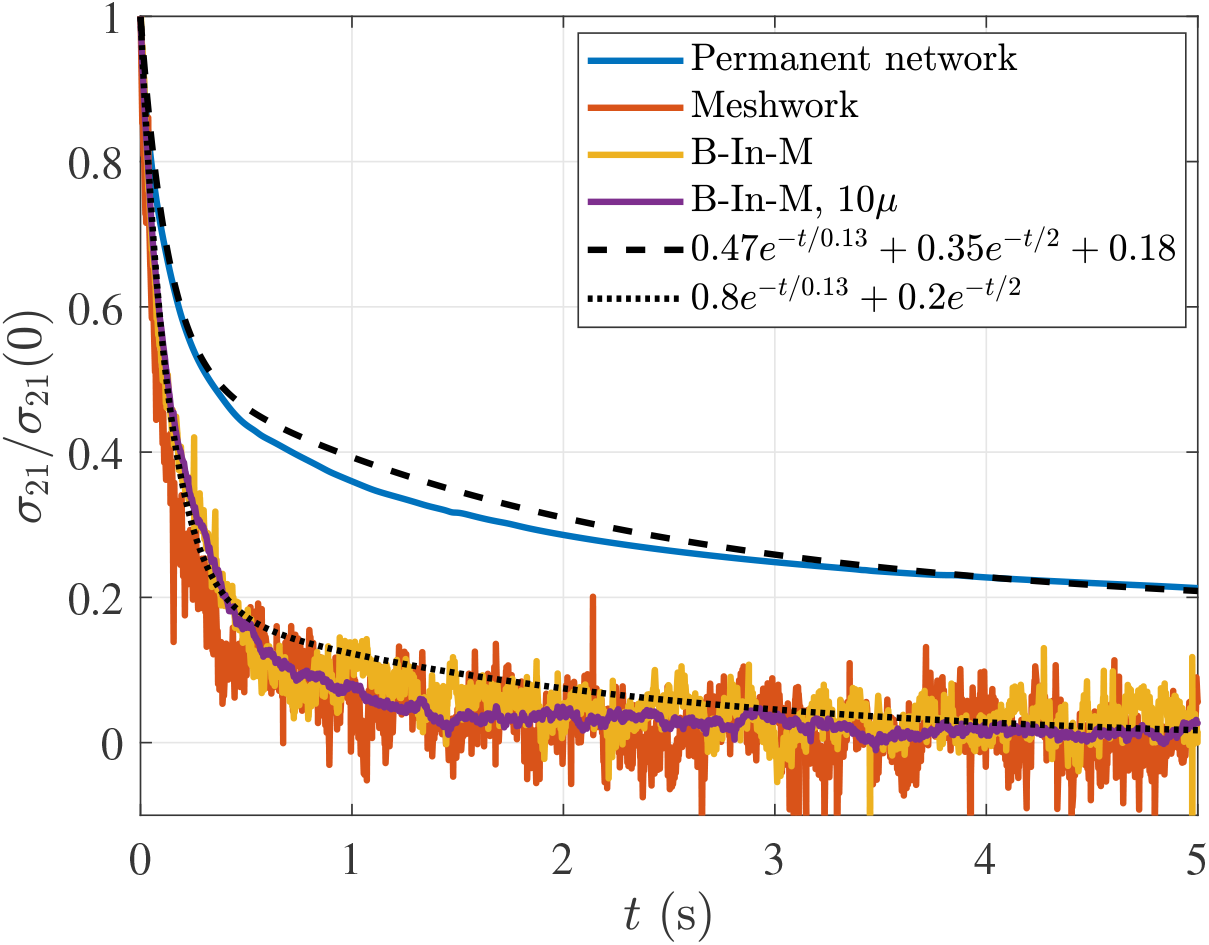
Normalized stress profiles over time in the stress relaxation test. We consider four different systems: a permanent network (blue), for which the stress relaxes to a nonzero value (*g*_0_ ≈ 0.07 Pa after accounting for normalization), and three dynamic networks, for which the stress relaxes to a value of zero (orange is the homogeneous meshwork, yellow the B-In-M morphology, and purple the B-In-M morphology with ten times larger viscosity). To illustrate the point that there are multiple intrinsic relaxation timescales in the system, we show a double-exponential curve which approximately matches the decay of stress for both permanent and transient CLs.

Figure 4 gives us two pieces of information that are important for our analysis going forward. First, we observe that statically-linked networks can store elastic energy, since *g*_0_ := *σ*_21_(*t* → ∞) > 0, while all of the energy dissipates in transiently-linked networks. Second, both permanent and dynamic networks have multiple relaxation timescales, as most of the stress (≈ 80% for dynamic networks) relaxes over the first 0.2 s or so, with the remaining part taking seconds to relax. In Fig. 4, we show a two-timescale decay curve that approximately, but not exactly, matches the decay of the stress in both statically- and transiently-linked networks. Given that one of the time constants is on the order 0.1 s and the other is on the order 1 s, Fig. 4 demonstrates that both slow and fast timescales exist in cross-linked networks. Our rheological experiments will give more precise estimates for the individual timescales, but it is worth noting that the dissipation of *all* of the stress over a 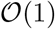 timescale implies that the longest timescale in the system cannot be much longer than five seconds. This is not surprising since the filament turnover time is of this order, but is in sharp contrast to modeling studies without turnover, where stress takes hundreds of seconds to dissipate [66, Fig. 4(a)].

One observation we can make from Fig. 4 is that the long-time behavior is similar for all dynamic networks, so the long timescale behavior is roughly independent of viscosity. We made this latter observation once before when we saw that a similar turnover time can be used to generate a B-In-M morphology for larger viscosity systems. We will look at the short-timescale behavior more closely in our rheological tests, which we discuss next.

### 3.3 Viscoelastic moduli

In Fig. 5, we plot the elastic and viscous moduli as a function of frequency for the five different systems defined in Table 2. We also include the data for a permanent, interconnected network of the type considered in [57], which we show with and without the constant elastic element *g*_0_ = *G*″(*ω* → 0) determined in Fig. 4. For the system with ten times larger viscosity, we show the data with time/frequency rescaled by a factor of ten, so that the value for *ω* = 1 Hz is mapped to *ω* = 10 Hz. These data are for the local drag model, since it is faster to simulate; we will consider the effect of nonlocal hydrodynamics in Section 3.7.

**Figure 5:**
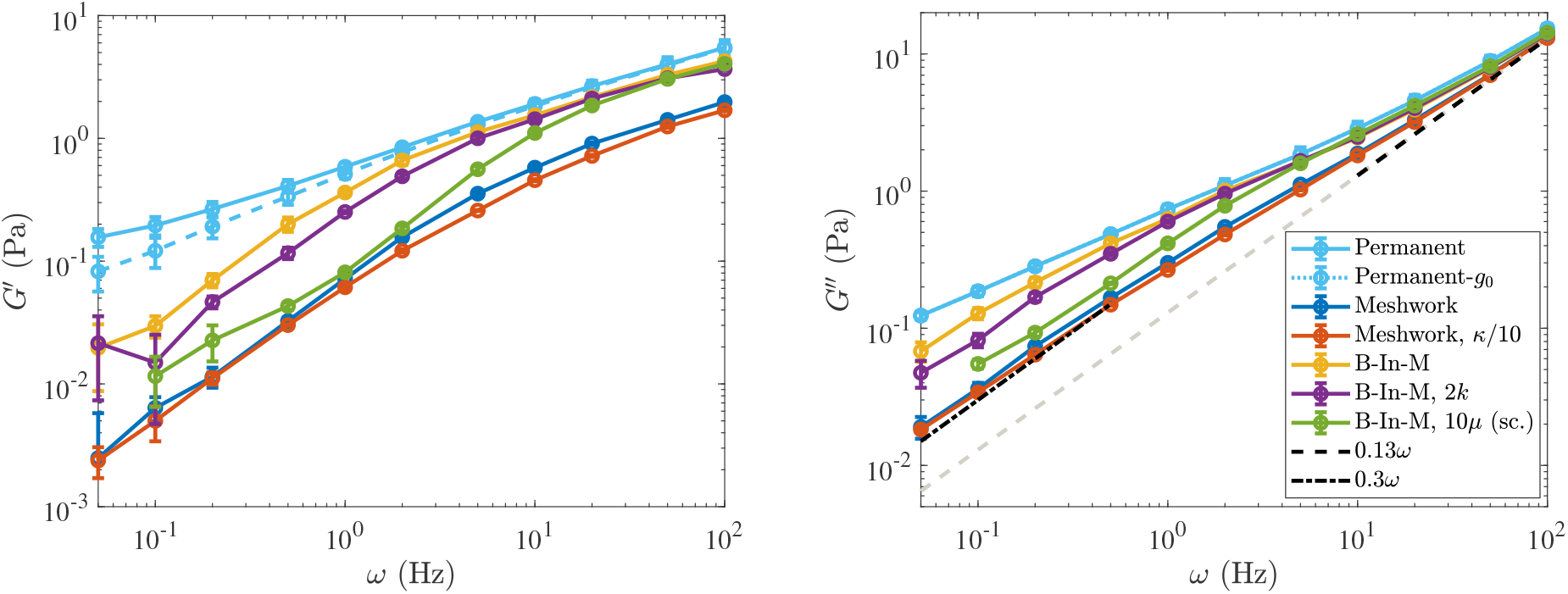
Elastic (left) and viscous (right) modulus for the five systems described in Tables 1 and 2 (see Table 2 for what parameters are varied in each system). For the B-In-M morphology with ten times the viscosity (*μ* =1 Pa·s), we show the data with time rescaled by a factor of ten. We also show the results for permanently-cross-linked networks, and for the elastic modulus the dashed light blue curve is the remaining elastic modulus when *g*_0_ ≈ 0.07 Pa (measured in Fig. 4) is subtracted off.

Examining the data, we make the following observations:

- On very short timescales (shorter than 0.05 s, or frequencies larger than 20 Hz), the viscous moduli all approach the viscous-fluid scaling of ≈ (0.13 Pa·s) *ω*. The timescale on which this occurs is directly proportional to the viscosity, as the rescaled *G*″ curve for the larger viscosity system makes the transition at the same time as the other B-In-M curves. In addition, the elastic moduli for B-In-M networks are about twice as large as those of homogeneous meshwork curves at high frequency, so the elastic modulus at short times is proportional to the link density.
- On very long timescales (several seconds, low frequencies), the viscous modulus again scales linearly with frequency, but with a larger slope (*G*″ ≈ (0.3 Pa·s) *ω*), which indicates that the links have become viscous on those timescales. The elastic modulus is much smaller than the viscous modulus on these timescales, and both of the moduli scale nonlinearly with the link density, as B-In-M morphologies have moduli about 4 times as large as homogeneous meshworks.
- There is an intermediate timescale of about 1 second on which the moduli for the B-In-M meshwork (yellow) diverge from those for the system with twice the binding/unbinding rate (purple), and on which both of these curves diverge from the light blue permanent network curve. In particular, for *ω* ≤ 2 Hz the elastic and viscous modulus are both smaller for the system with faster link turnover. This timescale is related to (in fact, it is exactly equal to) 1/*k*_off_, the timescale on which individual links bind and unbind. The rescaled curve with larger viscosity also diverges from the other B-In-M curves on this intermediate timescale, which indicates that a timescale has been introduced (by the dynamic linking) that does *not* scale with viscosity.

We will explore each of these fast, slow, and intermediate timescales in subsequent sections. The picture we sketch is of a rigid network for timescales shorter than the fastest intrinsic timescale *τ*_1_ ~ 0.1 s; on these timescales the links are predominantly elastic and the viscous modulus is the same as if the fibers were in a fluid without CLs. On timescales longer than *τ*_1_ but shorter than the intermediate timescale *τ*_2_ ~ 1 s, the links still appear permanent, but they provide additional viscosity by deforming the network to its steady state in response to flow deformations. On timescales longer than *τ*_2_, but shorter than the longest timescale *τ*_3_ ~ 5 s, the links come off before they are able to fully respond to the deformations induced by the background flow, and the slope of the elastic and viscous modulus changes relative to the one observed in permanent networks. Finally, on timescales longer than *τ*_3_, the network is almost completely viscous, and the viscosity is controlled by the morphology.

### 3.4 Short timescales: elastic links and viscous fibers

Our hypothesis is that the shortest timescale in the system, *τ*_1_, dictates when the links make a negligible contribution to the viscous modulus. For timescales *τ* < *τ*_1_, the network is essentially frozen: the fibers are rigid, the links are static, and the fibers and links contribute exclusively to the viscous and elastic modulus, respectively. Specifically, as *ω* → ∞, we expect that the links will make a constant contribution to *G*″ and that *G*″ will scale like *η*_0_*ω*, i.e., as a viscous fluid, with the viscous coefficient *η*_0_ independent of the number of links in the system. In Fig. 5, we show that *η*_0_ ≈ 0.13 Pa·s if *ω* is given in Hz. We have confirmed that this scaling continues for frequencies all the way up to 1000 Hz.

To verify that the viscous modulus at short timescales is dominated by the fibers only (independent of the number and nature of the links), we compare to the theoretical shear viscosity enhancement 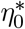 for an isotropic suspension of rigid cylindrical fibers, where the mobility is computed by the local drag approximation (first line in (8)). For dilute suspensions, the enhanced viscosity is given to order (log *ϵ*)^−2^ in terms of the fiber density *f* and aspect ratio *ϵ* as [7, 77, 53]

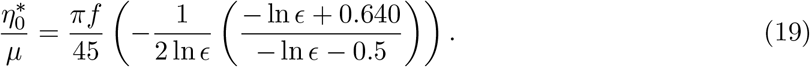

For our parameters (*ϵ* = 0.004, 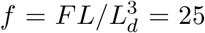), we obtain 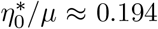 for cylindrical fibers.^3^ Recalling that *G*″ = 2*πωη*_0_ for a viscous fluid, our scaling in Fig. 5 shows that we obtain a similar value of *η*_0_/*μ* = 0.13/(2*π* × 0.1) ≈ 0.21.

It is important to separate the viscous contribution from the *links*, which approaches zero at short timescales, from that due to the *fibers*, which is infinite as *ω* → ∞. For this reason, we define

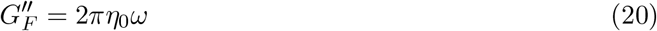

as the viscous modulus coming from the fibers. In Fig. 6(a), we examine the behavior of the moduli due to the links, *G*′ and 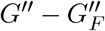, rescaled by the average number of links in the system 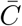. After rescaling time in the system with larger viscosity, we see that the data for *ω* > 10 Hz can be collapsed onto a single curve for both the elastic and viscous modulus. Thus on timescales shorter than *τ*_1_ ~ 0.1 s, each link behaves as an elastic element with strength independent of morphology. Note that the elastic modulus is more than 2 – 3 times larger than the viscous modulus (coming from the links) at high frequencies.

**Figure 6:**
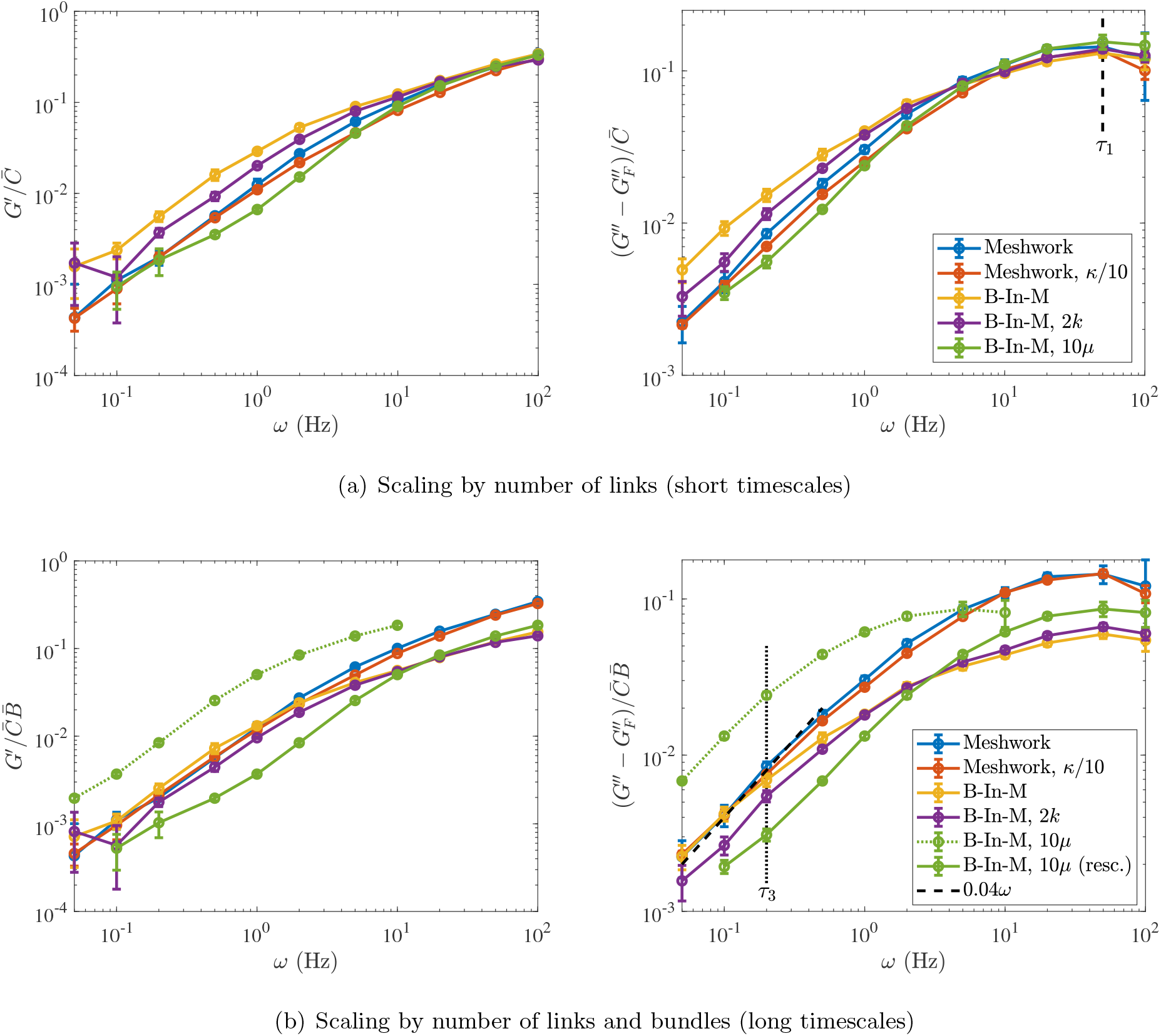
Elastic (left) and viscous (right) modulus *due to the CLs* for the five systems described in Tables 1 and 2 (see Table 2 for what parameters are varied in each system). To obtain a viscous modulus due to the links alone, we subtract the component due to the fibers 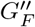 defined in (20). (a) We normalize by the link density 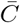 and see that the curves all match at short timescales on the order *τ*_1_ ≈ 0.02 s (after we also rescale time by viscosity). (b) We normalize by the link density multiplied by the bundle density. For the system with ten times larger viscosity, we also include the raw data (not rescaled) as a dotted green line. In the viscous modulus plot on the right, we show a linear slope as a dashed black line and define *τ*_3_ ≈ 5 s as the end of the low-frequency linear regime in *G*″. This timescale is roughly independent of viscosity.

### 3.5 Long timescales: viscosity depends on morphology and link unbinding rate

Let us now transition to the opposite limit of long timescales. At long timescales, Fig. 6(a) shows that each system is about five times more viscous than elastic (e.g., compare the values at *ω* = 0.2 Hz), and that normalizing by 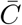 does not place the curves on top of each other. We say that these “long timescales” are longer than *τ*_3_, the longest intrinsic timescale in the system, or the timescale on which the network totally “remodels” itself. Our goal in this section is two-fold: first, we would like to find a new rescaling that better matches the curves at low frequencies, and second, we would like to estimate *τ*_3_, which we do by finding the regime in which *G*″ is a linear function of *ω* for each system.

Figure 6(b) shows our attempt to accomplish both of these goals. For both the viscous and elastic modulus, we rescale by the mean link density 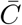 multiplied by the mean number of bundles 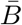, the idea being that bundles make a stronger contribution to the moduli at low frequency. We see a better match at low frequencies than in Fig. 6(a), which implies that bundles have a stronger effect on the mechanical behavior at long timescales than at short ones. In particular, the homogeneous meshworks (blue and orange) and B-In-M (yellow) curves all scale onto the same curve for frequencies *ω* ≤ *ω*_3_ ≈ 0.2 Hz = 1/*τ*_3_, but the curves with larger viscosity (green) and faster link (un)binding (purple) do not scale as well. In both of these cases, there is a change in a timescale shorter than *τ*_3_, as we discuss in the next section.

To estimate *τ*_3_, we determine the regime where the viscous modulus *G*″ scales *linearly* with *ω* and equate *τ*_3_ with the end of the linear region. From Fig. 6(b), we see that *τ*_3_ ≈ 5 seconds, and that the linear regime endures longer for homogeneous meshworks, which implies that they start becoming purely viscous at shorter timescales than systems with bundles (or, equivalently, these networks “remodel” themselves faster).

To understand exactly how much viscosity is being provided by the links and bundles in the different systems, we use the slope in Fig. 6(b) to obtain

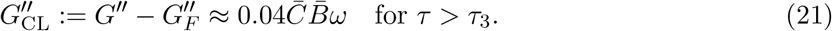

Since 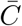 has units links per fiber and 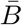 has units bundles per *μ*m^3^, 0.04 in this equation has units Pa·s/*μ*m^3^. To extract a viscosity, we write 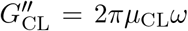, so that in the B-In-M system with 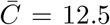 and 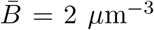 we obtain *μ*_CL_/*μ* = (0.04 × 12.5 × 2)/(2*π* × 0.1) = 1.6. This is eight times more viscosity than the fibers alone (0.2) and 1.6 times the background fluid viscosity. For the homogeneous meshwork with 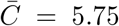 and 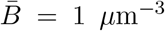, the additional viscosity is *μ*_CL_/*μ* = (0.04 × 5.75 × 1)/(2π × 0.1) = 0.4. This is twice the viscosity of the fibers but 2.5 times smaller than that of the background fluid. Comparing the two morphologies confirms some of the prior intuition that bundles embedded in meshworks provide stronger resistance to flow than homogeneous meshworks [85].

### 3.6 Intermediate timescales

So far, we have addressed the two extreme limits of the suspension behavior. Over short timescales, *τ* < *τ*_1_, the entire network is rigid, the CLs are purely elastic, and the additional viscosity is the same as that due to the fibers alone. Over long timescales, *τ* > *τ*_3_, all of the energy in the network is dissipated. Bundles break up, there is no elastic modulus, and the morphology of the network, and the amount by which links can deform it, determines the viscous behavior.

There is also an intermediate timescale *τ*_2_ at which *some of* the links, but not the entire network, start to behave viscously. While we expect *τ*_2_ ~ 1/*k*_off_, this could change based on the network parameters, as the same value of *k*_off_ acts differently depending on the speed of network deformation. To better understand the timescale *τ*_2_, we compare the elastic modulus with dynamic links to the one for statically-linked networks (by starting from the same equilibrium structure and keeping the links fixed). Similar to [57], we do this for five seconds or five cycles, and measure the modulus over the last three cycles.

Figure 7(a) shows the results for our B-In-M structures. The solid lines show the elastic modulus with dynamic linking, and the dotted lines show the elastic modulus with permanent linking. The timescale *τ*_2_ is when the dynamic linking elastic modulus diverges significantly from the permanent link modulus, so that the links come off before they can provide their maximum elastic response. From Fig. 7(a), we see that *τ*_2_ is obviously smaller when we increase the rate of link turnover (the purple curve diverges from the dotted yellow one faster than the solid yellow one does); this is expected from *τ*_2_ ~ 1/*k*_off_, which is 1 s for the yellow curve and 0.5 s for the purple one.

**Figure 7:**
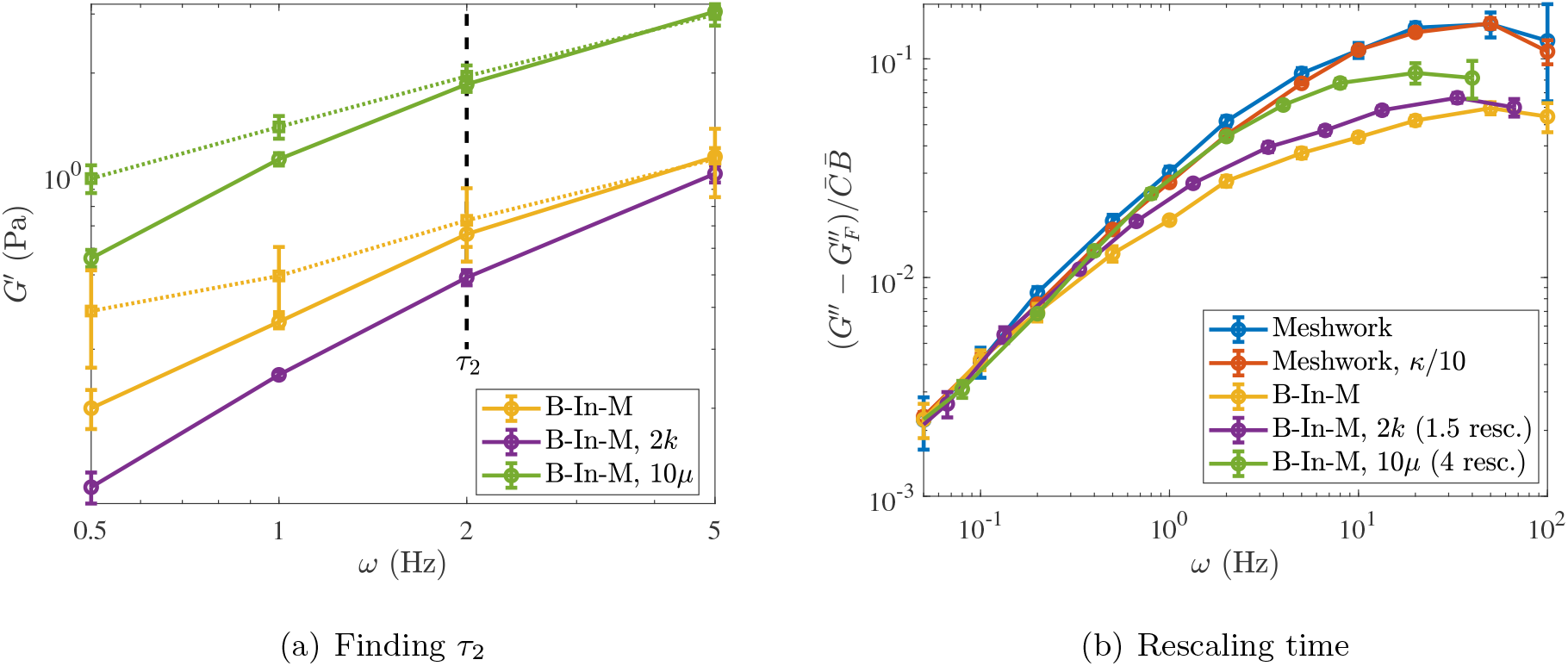
(a) Elastic modulus at medium frequencies. For each network type (indicated in the legend), we compare the solid line, which has the elastic modulus with dynamic links, to the dotted line, which is the elastic modulus with permanent links. The intermediate timescale *τ*_2_ ≈ 0.5 s is the inverse of the frequency where the data start to diverge. We consider only bundle-in-mesh morphologies using our standard parameters (yellow), twice the link turnover rate (purple, but the static link reference is still the dotted yellow), and ten times the viscosity (green). (b) Rescaling time to get all of the viscous modulus curves on the same plot. This is the same plot as Fig. 6(b) (right panel), but now we rescale the time for the larger viscosity green curve by a factor of 4 instead of 10 (*ω* → 4*ω*), and we rescale the time for the faster link turnover purple curve by 1.5 (*ω* → *ω*/1.5). This demonstrates that the data do not scale simply with the parameters at medium and low frequencies, when multiple timescales are involved.

Morphology has no impact on the timescale *τ*_2_, as we obtain the same characteristic time *τ*_2_ ≈ 1 s for the homogeneous meshwork (not shown, because the results are indistinguishable) as the B-In-M morphology. Thus the morphology only appears to have a strong influence on the longest timescale *τ*_3_.

It is harder to determine the effect of viscosity on *τ*_2_. On the one hand, Fig. 7(a) shows the true data for *G*″ vs. *ω* (without scaling time by *μ*), so it appears that the dynamic curve diverges from the permanent one at about the same timescale regardless of viscosity. On the other hand, for a fixed *k*_off_ the difference between the solid and dashed curves at *ω* = 1 Hz is larger for larger viscosity (green). Thus the timescale *τ*_2_ depends on the interaction of dynamic linking and deformation in a nontrivial way.

To illustrate this complicated dependency, we show that different rescalings are required when the viscosity (dynamics) or link turnover rates change from the base parameters. We recall from Fig. 6(b) that the larger viscosity (green) and faster link turnover (purple) B-In-M systems do not follow the same normalization as the other systems. Figure 7(b) shows that scaling time by a factor of 4 (and not the expected 10) puts the green larger viscosity curve onto the rest of the curves at low frequencies, while scaling time by a factor of 1.5 (and not the expected 2) puts the curve with twice the link turnover rate on top of the others as well. The reason for these particular scalings is not obvious to us, other than that they fall somewhere between the expected scaling and unity.

### 3.7 Role of nonlocal hydrodynamic interactions

We have seen that the network behaves very differently on short timescales, where each link behaves elastically independent of the network morphology, than on long timescales, where the system is purely viscous and the morphology exerts a significant influence on the resistance to flow. In this section, we show that nonlocal hydrodynamic interactions significantly decrease the elastic and viscous moduli in B-In-M morphologies. We show that the decrease is most significant at low frequencies, i.e., the frequencies where we already know morphology has a strong influence on the behavior. We explain the decrease in two ways: first, nonlocal hydrodynamic interactions slow down the bundling process, causing less bundles to form for a fixed turnover time, and second, they create entrainment flows that reduce the stress inside of the bundles that do form.

We consider only the two characteristic morphologies in this section without varying any other parameters from those given in Table 1. For each of the morphologies, we compute the viscoelastic moduli with nonlocal hydrodynamics, then compute them again using local drag and intra-fiber hydrodynamics. In Fig. 8, we plot the error/fraction of the moduli recovered with the various mobility operators. For a homogeneous meshwork system (blue lines), we see that the elastic modulus is the same within 10 – 20% whether we use local drag, intra-fiber, or full hydrodynamics. At low frequencies, the viscous modulus is also the same within 10% regardless of the hydrodynamic model used, but there is an obvious error in the viscous modulus when the local drag model is used at high frequencies. This discrepancy of about 20% is also present in permanent networks [57, Fig. 9(b)] and in B-In-M morphologies, and in all cases it can be mostly recovered by adding only *intra-fiber* hydrodynamics to the mobility. The 20% difference between local drag and intra-fiber hydrodynamics is similar to the theoretical estimate discussed in footnote 3. There appears to be a small increase (at most 5%) in the viscous modulus when we switch from intra-fiber hydrodynamics to fully nonlocal hydrodynamics, which would be more significant at smaller mesh sizes [77].

**Figure 8:**
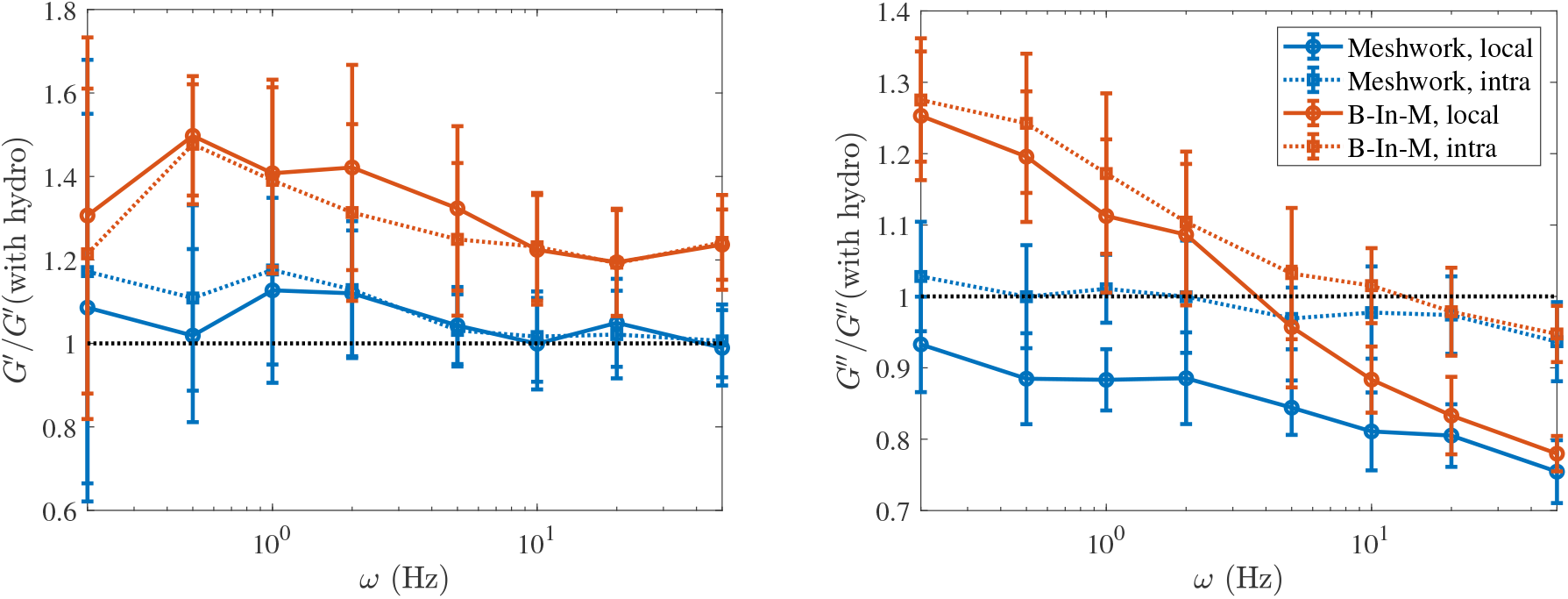
Proportion of the (left) elastic and (right) viscous modulus that is recovered using various mobility approximations. We compute the modulus using full hydrodynamics, then plot the fraction of it recovered using local drag or intra-fiber hydrodynamics. We show the results for the homogeneous meshwork in blue and the B-In-M morphology in orange. For each line color, a solid line shows the results for local drag and a dotted line shows the results for intra-fiber hydrodynamics. Intra-fiber hydrodynamics cannot explain the deviations in the elastic and viscous modulus at low frequency for the B-In-M geometry.

**Figure 9:**
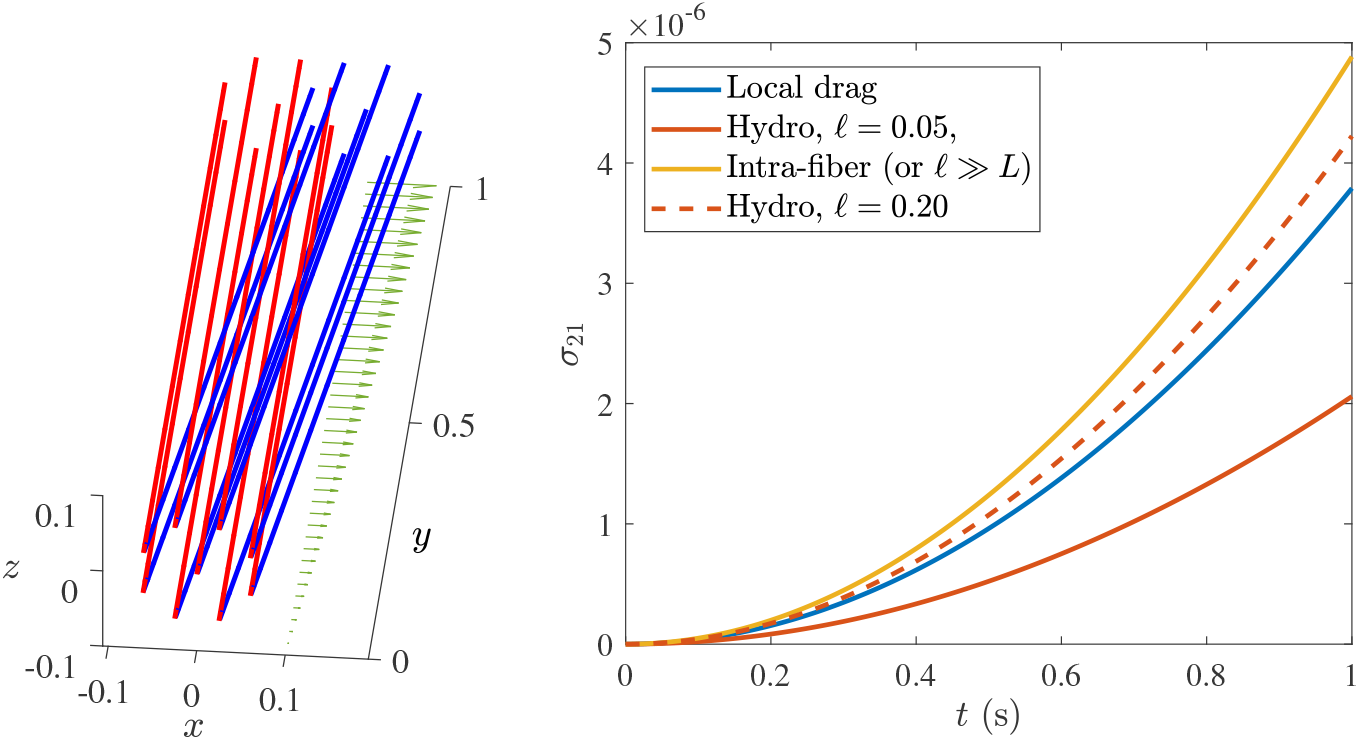
Reduction of stress in bundles explains smaller moduli with hydrodynamics. (Left) We manufacture a bundle geometry without CLs by placing nine fibers of length *L* =1 *μ*m (red) in and around an octagon with side length ℓ and straining with constant rate 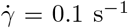 until *t* = 1 s (blue). (Right) The resulting stress evolution for different hydrodynamic models. The local drag (blue) and intra-fiber (yellow) results are independent of ℓ, while the stress for full hydrodynamics (orange) depends strongly on ℓ. For ℓ = 0.05 *μ*m (solid orange, the simulation parameters), there is a significant decrease in stress which comes from the entrainment of the fibers in each other’s flow fields. For ℓ = 0.20 *μ*m (dashed orange), the decrease is minimal; note that for full hydrodynamics with ℓ ≫ *L* we would recover the “intra-fiber” curve.

The more interesting changes with nonlocal hydrodynamics come at low frequencies in the B-In-M geometries. Unlike with the homogeneous meshwork, the local drag model makes about a 50% overestimation in the elastic modulus and a 30% overshoot in the viscous modulus at low frequencies for B-In-M morphologies, with the largest change coming at *ω* = 0.5 Hz, or a timescale of 2 seconds. We have previously seen that morphology has a strong influence on the moduli at these timescales, so we explore the impact of nonlocal hydrodynamics on morphology next.

#### 3.7.1 Rescaling of time cannot explain long-timescale moduli

The first possible explanation for the decrease in *G*″ and *G*″ with nonlocal hydrodynamics at low frequencies is a change in the network structure. In this section, we show that, while there are less bundles and links in the dynamic steady state with full (intra- and inter-fiber) hydrodynamics for a given turnover time, this by itself cannot entirely explain the decrease in the moduli.

To do this, we focus on the frequency where full hydrodynamics matters the most as a percentage. For *ω* = 0.5 Hz, we measure the moduli with intra-fiber hydrodynamics as 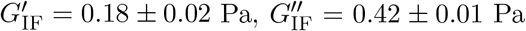, while the moduli with full hydrodynamics are 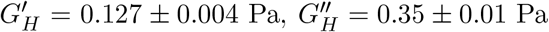. As shown in Table 4, the steady state link and bundle density are 12.3 and 2.2 *μ*m_^3^, respectively, with intra-fiber hydrodynamics, while with full hydrodynamics they are are 11.7 and 2.0 *μ*m^−3^. Thus at its dynamic steady state, the network has on average fewer links and fewer bundles when we simulate with full hydrodynamics than when we simulate with intra-fiber hydrodynamics. This is because disturbance flows in bundling fibers (fibers moving towards each other) *oppose* the direction of motion, which slows down the bundling process. Thus when we fix a turnover time, there are on average fewer bundles (and fewer links) when the bundling process is slower.

**Table 4.**
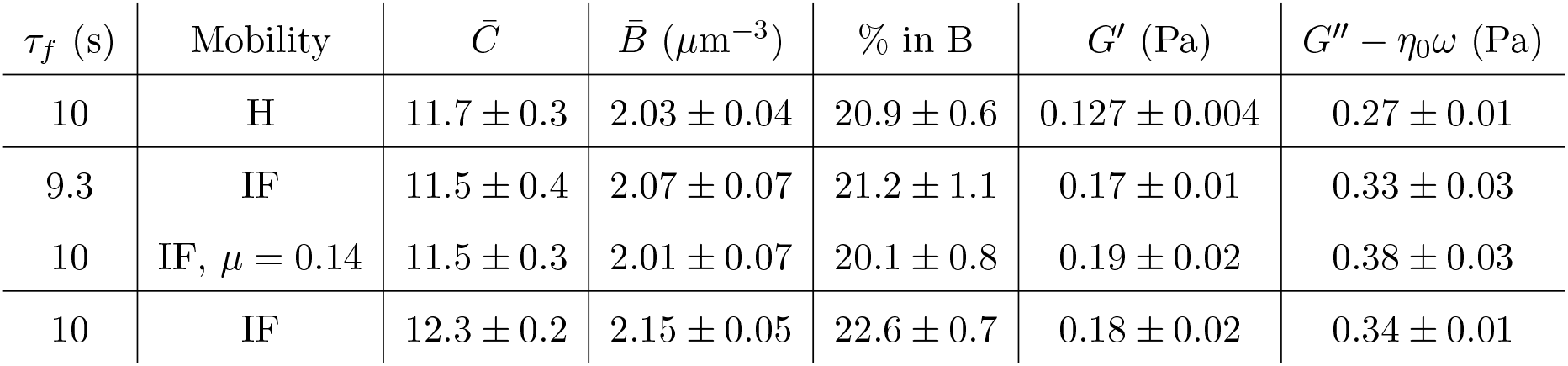
Statistics for system with *ω* = 0.5 Hz and varying turnover times and mobility models (IF stands for intra-fiber hydrodynamics where from Table 5 we obtain *η*_0_/*μ* = 1.6, and H stands for full, nonlocal hydrodynamics, where *η*_0_/*μ* = 1.7). In the second and third row, we attempt to tune the turnover time or viscosity so that the steady state morphology with intra-fiber hydrodynamics matches the steady state morphology with hydrodynamics and *τ_f_* = 10 s. Even after doing this, the elastic and viscous modulus are still significantly higher with the intra-fiber mobility.

To compensate for the changes in structure, we drop the turnover time or increase the viscosity for intra-fiber hydrodynamics simulations so that the steady state morphology matches the morphology with hydrodynamics as closely as possible. We then measure the moduli for these new steady states. In Table 4, we show our attempt to match the statistics for intra-fiber hydrodynamics with smaller turnover times with those for hydrodynamics with *τ_f_* = 10 s. Comparing *τ_f_* = 10 s with full hydrodynamics with *τ_f_* = 9.3 s with intra-fiber (IF) hydrodynamics, we see the IF simulations have larger moduli, even when we closely match the steady state link density 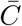 and bundle density 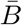. The same is true when we attempt to rescale time by increasing the background fluid viscosity (second to last line in Table 4), as in that case the larger values of *G*′ and *G*″ persist despite a decrease in link and bundle density. So nonlocal hydrodynamics must both change the bundle morphology and lower the stress for a fixed morphology.

#### 3.7.2 Bundled fibers: when flow reduces stress

The reason for the decrease in moduli with inter-fiber hydrodynamics for a fixed morphology has to do with the flows inside a bundle. If the fibers are packed tightly within a bundle, then we expect their disturbance flows to have a strong influence on each other. When a fiber in a bundle moves with the bulk fluid, the disturbance flow it creates is in the same direction as the bulk motion, so the other fibers in the bundle will naturally move as well. This effect, which is not present when we use local drag or intra-fiber mobility, explains the reduction in stress for a fixed rate of strain.

Figure 9 establishes this more rigorously. We consider a set of nine fibers with *L* = 1 *μ*m *without* cross-links. The fibers are arranged in a regular octagon with side length ℓ with another fiber centered at the origin. We apply a constant straining flow with strength 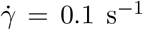 and measure the stress over the first second in Fig. 9. The solid orange line shows the stress with full hydrodynamics and the simulation parameter of ℓ = 0.05 *μ*m, so that the fibers are very close. In this case, we see that the stress with full hydrodynamics is slightly more than half of that with local drag (which is independent of ℓ). Similar to our simulation result for cross-linked gels, the stress decrease is *not* explained by intra-fiber hydrodynamics, which increases the stress from local drag by the theoretically predicted 20 – 25%. In fact, as discussed in footnote 3 and [57, Sec. 6.3.2], disturbance flows from intra-fiber hydrodynamics (which try to stretch the fiber) make the constraint forces on each fiber larger, which causes an increase in stress. Thus the decrease in stress with nonlocal hydrodynamics comes from the nonlocal flows induced by the fibers on each other. We demonstrate this in Fig. 9 by spacing the fibers farther apart (ℓ = 0.2 *μ*m) and showing that the stress with full hydrodynamics increases towards the intra-fiber curve.

### 3.8 A continuum framework: the generalized Maxwell model

In order to enable whole-cell modeling, it is important to introduce a continuum model for the passively cross-linked actin gel. We are motivated by the behavior of the network: at short timescales, it scales as a viscous fluid with a constant elastic modulus (a dashpot in parallel with a spring), while at long timescales it is purely viscous. Combining these two, we introduce a generalized Maxwell model of the type shown in Fig. 10. We have the viscous dashpot of strength *η*_0_ in parallel with three Maxwell elements, each of which has a strength *g_i_* and an associated relaxation timescale *τ_i_*, on which the element goes from elastic (for timescales shorter than *τ_i_*) to viscous (longer than *τ_i_*). The total elastic and viscous (in excess of the solvent viscosity) modulus are given by sums of the viscous dashpot and each Maxwell element [63]

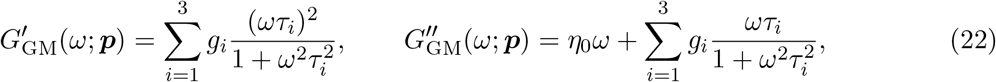

where we have denoted the 7 parameters by a vector ***p***. As discussed in [93, 58], we can obtain the fitting parameters (*g_i_, τ_i_, η*_0_) by maximizing the log-likelihood function

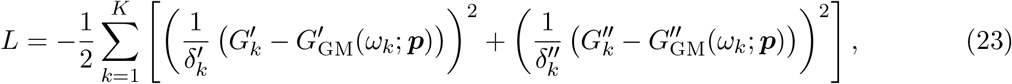

where 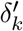 and 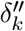 are the uncertainties in *G*′(*ω_k_*) and *G*″(*ω_k_*), respectively, and *K* is the number of frequencies studied. Weighting the observations by uncertainty will cause the fit to give more weight to more certain measurements. The uncertainty in the fit 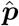 is estimated by the square root of the diagonal entries of the inverse of the Fisher information matrix,

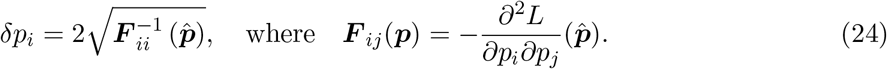

so that *p_i_* ± *δp_i_* is a 95% confidence interval for *p_i_*.

**Figure 10:**
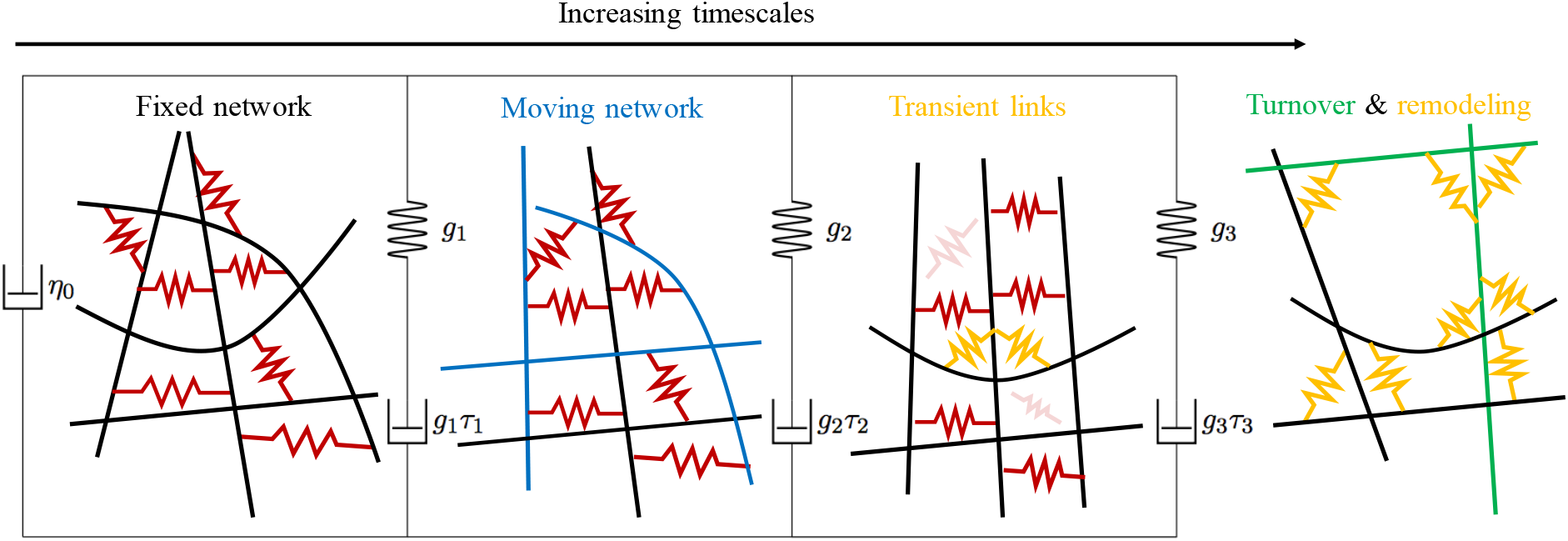
Our continuum model, informed by the timescales discussed in previous sections. We use three Maxwell elements with timescales *τ*_1_, *τ*_2_, and *τ*_3_, all in parallel with a viscous dashpot to describe the network. The viscous dashpot *η*_0_ represents the high frequency viscosity of the permanently cross-linked fiber suspension. The first Maxwell element has timescale *τ*_1_ ≈ 0.02 seconds associated with it, and represents the relaxation of the fibers to a transient elastic equilibrium (the networks before and after relaxation are shown to the left and right of this Maxwell element; the relaxing fibers are shown in blue); on this timescale, the links are effectively static. The second Maxwell element, with timescale *τ*_2_ ≈ 0.5 s, represents the unbinding of some links (shown more transparent than the others) and the appearance of new links (orange) – compare the networks to the left and right of this Maxwell element. The third Maxwell element with timescale *τ*_3_ ≈ 5 s represents network remodeling (compare the networks to the left and right of this element); for timescales larger than *τ*_3_, some of the fibers (shown in green) and links (orange) turn over and the network completely remodels from the initial state.

In Fig. S5(a), we verify that using three (as opposed to two or four) Maxwell elements is indeed the best choice for the bundle-embedded meshwork with *τ_f_* = 10 s and full hydrodynamics (the other systems are similar). As detailed in [9, 58], we determine this by increasing the number of Maxwell elements until the fit stops improving substantially and we start to overfit the data, leading to ill-conditioning. In addition to comparing the fit with three timescales to that with two or four, in Fig. S5(a) we also compare to a fit using a continuum of timescales, which is suggested by physical theories [10, 65] and used in prior studies [11]. In this approach, the discrete values of *g_i_* in (22) are replaced by a continuous function *g*(*τ*), which is integrated (instead of summed) on 0 ≤ *τ* ≤ *τ_max_*. If we take the common choice of *g*(*τ*) = *g*_0_*τ*^−*α*^ [55], we obtain

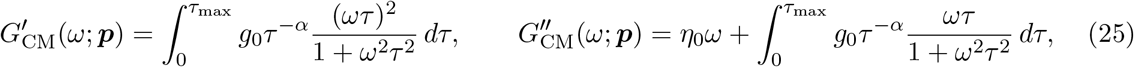

where (*g*_0_, *α*, *τ*_max_, *η*_0_) are the fitting parameters. In Fig. S5(a), we show that the fit to the data with this continuum spectrum of timescales is not substantially better than the three timescale generalized Maxwell fit. Our preference is the three timescale fit, since in this case we can assign physical to meaning to each of the timescales, as we do in Sections 3.4–3.6. Given that adding more discrete timescales leads to ill-conditioning, the continuum approach, where there are infinitely many timescales (and in fact infinitely many for choices *g*(*τ*)), is even more poorly conditioned, and in that case it is difficult to assign a physical meaning to the fitting parameters.

The three-timescale fit that we obtain for the B-In-M system with full hydrodynamics, along with the contribution of each of the separate Maxwell elements, is shown in Fig. 11. Admittedly, the elastic modulus data *G*′(*ω*) at small frequencies *do not* fit the generalized Maxwell model since they do not decay as *ω*^2^. This suggests that the material response is likely more complicated than just three Maxwell elements. At the same time, we have already shown that *G*″(*ω*) scales linearly with *ω* at small frequencies, and thus that the viscous modulus, which dominates the elastic modulus at low frequencies, does fit the generalized Maxwell assumption. Our choice is therefore to fit only *G*″, and not *G*′, for frequencies less than 0.1 Hz (10 s timescales).

**Figure 11:**
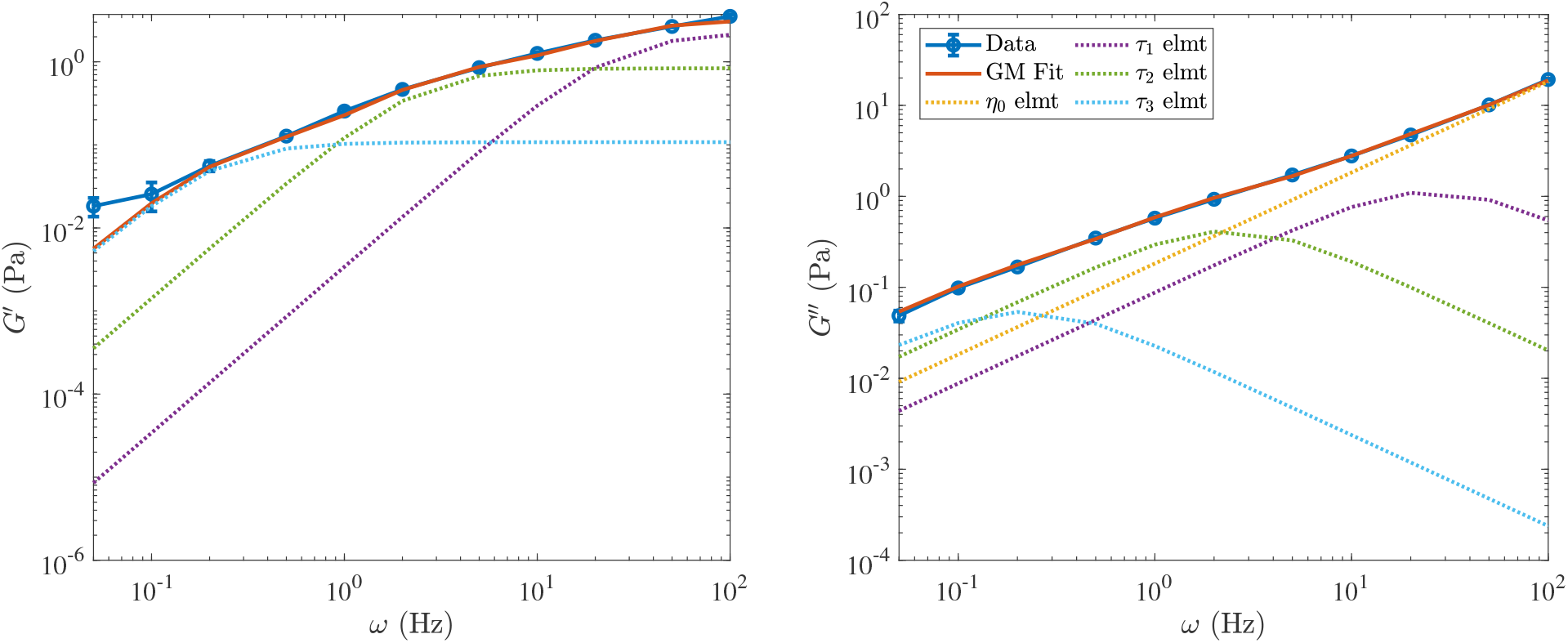
Fitting the model of Fig. 10 (the moduli (22)) to the data from the B-In-M system. The data are shown in blue, with the total fit shown in red. The dotted lines show the contribution of each element in Fig. 10 to the total fit. Here *τ*_1_ = 0.04 s is the fastest timescale shown in purple, *τ*_2_ = 0.41 s is the intermediate timescale shown in green, *τ*_3_ = 4.5 s is the longest timescale shown in light blue. The yellow dotted line shows the contribution of the pure viscous element.

Table 5 gives the fitting parameters for all of the systems we have considered with multiple options for the mobility. For almost all systems, we have *τ*_1_ ≈ 0.03 s, *τ*_2_ ≈ 0.3 s, and *τ*_3_ ≈ 3 s, which is in line with our estimates of these timescales in Figs. 6 and 7(a). The only exception is the system with higher viscosity; in this case we have already shown that the timescale *τ*_1_ scales with viscosity to become about 0.3 s, while the timescale *τ*_2_ remains about 0.5 s. The result is that *τ*_1_ and *τ*_2_ blend into a single timescale, and a two-timescale model is the best choice for the data, as we show in Fig. S5(b). Comparing B-In-M and meshwork geometries, we observe that the long timescale *τ*_3_ is either longer (comparing B-In-M and meshwork with full hydrodynamics) or its contribution in (22) is stronger (comparing B-In-M and meshwork with local drag), which is consistent with our observation in Section 3.5 that B-In-M geometries have longer remodeling times and a higher resistance to deformation. Finally, we notice how the viscosity *η*_0_ scales directly with *μ* and increases by ≈ 30% when we account for intra-fiber or nonlocal hydrodynamics, in accordance with our observations in Sections 3.4 and 3.7.

**Table 5.** Parameters in the three-timescale generalized Maxwell model shown in Fig. 10. Refer to Table 2 for a detailed description of each system. The uncertainty in each measurement is about 25%, with the exception of *η*_0_, which has an uncertainty of about 5%.

| System | Mobility | $(\tau_1, \tau_2, \tau_3)$ (s) | $(g_1, g_2, g_3)$ (Pa) | $\eta_0$ (Pa·s) |
| --- | --- | --- | --- | --- |
| Meshwork | LD | (0.032, 0.31, 4.2) | (1.56, 0.44, 0.03) | 0.13 |
| Meshwork, $\kappa/10$ | LD | (0.024, 0.15, 1.5) | (1.34, 0.45, 0.07) | 0.13 |
| Meshwork | Intra-fiber | (0.022, 0.17, 1.4) | (1.49, 0.59, 0.10) | 0.16 |
| Meshwork | Full hydro | (0.025, 0.21, 2.1) | (1.42, 0.54, 0.05) | 0.17 |
| B-In-M | LD | (0.026, 0.28, 2.7) | (2.91, 1.26, 0.30) | 0.13 |
| B-In-M | Intra-fiber | (0.030, 0.28, 2.4) | (2.77, 1.17, 0.29) | 0.16 |
| B-In-M | Full hydro | (0.039, 0.41, 4.5) | (2.26, 0.84, 0.11) | 0.18 |
| B-In-M, $2k$ | LD | (0.034, 0.31, 2.7) | (2.54, 1.10, 0.17) | 0.13 |
| B-In-M, $10\mu$ | LD | (0.49, —, 4.3) | (3.13, —, 0.38) | 1.33 |

## 4 Discussion

We experimented numerically with dynamic actin gels made of micron-long actin fibers at concentrations characteristic of in vitro experiments. In the simulations, the fibers turned over rapidly on a 5 to 10 second scale, similar to some experimental measurements [82, 27]. The gel was cross linked dynamically with flexible spring-like cross linkers (CLs) characterized by mechanical and kinetic parameters similar to those for *α*-actinin, a ubiquitous actin CL. We observed that gels with faster fiber turnover evolved into relatively homogeneous meshworks, while gels with slower turnovers morphed into bundle-in-mesh (B-in-M) networks with small actin bundles immersed into cross-linked meshworks. This result agrees with experimental observations [30].

Our simulations revealed three principal time scales characterizing the mechanics of dynamic cross-linked actin gels. On the fastest time scale, ~ 0.03 seconds, actin fibers relax viscously, locally, and rapidly to the transient elastic equilibrium generated by the network configuration. On intermediate timescales, ~ 0.5 seconds, the CLs turn over, generating new transient elastic equilibria, and finally on the slow time scale, ~ 3 – 5 seconds, the network undergoes global remodeling through a combination of filament and CL turnover. One of the most useful practical results of the simulations is that very complex cross-linked gel mechanics can be approximated with the generalized Maxwell mechanical circuit shown in Fig. 10, in which three Maxwell elements in parallel correspond to elastoviscous gel deformations on the three characteristic time scales, in addition to the effective viscosity of the actin fiber suspension in the background fluid.

Our simulations predict that at small frequencies (time scales longer than a few seconds), the effective elastic and viscous behaviors are controlled by the CL mechanics (as opposed to fiber mechanics). In this regime, an effectively viscous mechanical response to deformation dominates relatively weak elastic behavior. At low frequencies, the dominant viscous response originates from the CLs stretching and deforming the fibers (in addition to the significant viscosity of the background fluid); the fibers by themselves contribute little to the net viscosity. The mechanical moduli scale nonlinearly with the CL density, because the CLs within actin bundles respond to shear differently than the CLs in the actin mesh. Thus, the gel’s morphology exerts a significant influence on the actin resistance to flow at long timescales.

At high frequencies, or timescales shorter than one tenth of a second, we found that the effective elastic modulus of the network is higher than (but on the same order of magnitude as) the effective viscous modulus (once we removed the viscous scaling generated by the fibers). The elasticity simply originates from multiple elastic springs of CLs that can be considered static on the short time scale. The greatest contribution to the net viscosity is that of the background fluid, with the fiber suspension contributing about 25% additional viscosity. At moderate frequencies around 1 Hz (intermediate timescale, between 0.1 and 3 seconds), the mechanical behavior of the crosslinked network is complex, combining comparable contributions of all mechanical factors described for low and high frequencies.

Our model allowed us, for the first time, to estimate quantitatively the role of nonlocal interfiber hydrodynamic interactions on the actin gel rheology. We demonstrated that, while these interactions have little effect in a homogeneous cross-linked actin meshwork, they significantly decrease the elastic and viscous moduli of heterogeneous B-In-M gels. We showed that this decrease is most significant at low frequencies, i.e., on timescales of a few seconds, when the network morphology has a strong influence on its mechanical behavior. At these frequencies, in the presence of bundles, the hydrodynamic interactions cause 30–50% downward corrections to the viscoelastic moduli. Two factors are responsible for this effect: first, hydrodynamic interactions slow down the bundling process, so fewer bundles form, and second, nonlocal hydrodynamics creates entrainment flows that reduce the stress inside of the bundles that do form.

While almost all of our simulations fell in the linear viscoelastic regime, we did observe a strain hardening effect, with the network resistance increasing beyond the linear response for higher-amplitude deformations. This effect has been observed before experimentally [22], which has prompted a variety of explanations and computational studies. Both Mulla et al. [65] and Kim et al. [40], for instance, propose that strain hardening is due to the mechanics of individual filaments, which stiffen nonlinearly in response to applied stress. Because our goal was to measure the viscoelastic moduli of the networks in their dynamic steady state, we did not systematically explore the viscoelastic behavior at large strains or perform measurements at zero frequency. That said, our simulations do offer two interesting insights: first, the mechanical behavior becomes nonlinear at just 10% or greater strains, which is a typical result for in vitro networks [79], and second, there are more bundles (which are more resistive to strain) at larger strain rates, and so at least one reason for the nonlinearity is the mechanosensitive character of the actin gel morphology. Our conclusion is identical to the modeling study [5], which found that strain combined with dynamic cross-linking induces signficant morphological changes in the network, leading to strain hardening [5, Fig. 3]. In fact, it was recently shown that actin filament mobility is a necessary condition for strain hardening and mechanical hysteresis [75].

In the linear response regime, our results are in line with most of the prior experimental and modeling studies. We found viscoelastic moduli on the order 0.1 – 10 Pa [23, 38], with the links becoming viscous on long timescales and elastic on short ones [11]. Our measurements showed that decreasing the fiber bending stiffness decreased the viscoelastic moduli, but by a slower than linear rate [42] (in our study, decreasing *κ* by a factor of ten reduced the moduli by at most 30%). We also found that the elastic stress in the gel (from the CLs exerting force on the fibers) is much larger than the viscous drag stress over long times, which agrees with a previous computational study [40].

We failed, however, to observe a local maximum or minimum in the viscous modulus at intermediate frequencies, as was reported in [48, 90]. One explanation for this is a difference in parameters. In the computational study [90], the unloaded CL unbinding rate is 0.1 s^−1^, and the CL stiffness is 10 pN/nm (10,000 pN/*μ*m). In this parameter regime, the CLs *always* make the most of their mechanical contribution to the network mechanics while bound, since they cross-link the fibers for seconds and are extremely stiff. Indeed, [90, Fig. 6] shows that the peak in *G*″, when the timescale of the driving frequency matches the link unbinding rate, becomes less sharp and almost disappears when *k*_off_ increases to the order of one second, as it is in our study here. This suggests that the local maximum is only observed when the relaxation of the network is many orders of magnitude faster than the link binding rate. This issue clearly requires more investigation, and in the future we will explore how using a force-dependent unbinding rate, as is done in [90], impacts the results, and whether this assumption is responsible for the peak.

What all modeling studies, including ours, suffer from is that only few features of the actin gel are simulated, with many factors, forces, and processes either ignored or approximated crudely. There is, of course, a good reason for this: even a very limited caricature of the gel exhibits complex dynamics; the fact that many experimental studies paint very different pictures of the gel mechanics likely stems from the same biological complexity. We propose that to understand the full complexity, one has to keep adding dynamic processes to previous simpler benchmark models and examine the changes that new dynamics bring. Thus, our next step will be to add thermal forces responsible for random bending of the fibers. For a one micron long fiber with persistence length close to 20 microns, the thermal bending causes lateral fluctuations of no more than 30 to 40 nm, which is significantly less than the mesh size of the networks we investigated, so we do not expect that our results will change drastically. Indeed, a previous computational study in permanent networks showed that thermal fluctuations only impact the mechanical moduli when the average distance between cross links is greater than the persistence length of the fibers [42, Fig. 7(c)], which is a regime we are far outside of here.

Despite this speculation, the fact remains that our study was unable to reproduce a number of experimental observations in transiently cross-linked actin networks that have been traced to the underlying thermal fluctuations of the filaments. For instance, we report a viscous modulus that decays as for a viscous fluid (*G*″ ~ *ω* at low frequencies), whereas a number of recent experimental, modeling, and theoretical studies [66, 10, 95, 96] report an *ω*^1/2^ dependence of both moduli at low frequencies. It is important to note, however, that fiber turnover, which was not accounted for in these previous studies, changes the behavior on timescales larger than the turnover time, as does our use of shorter filaments relative to [66]. In the high-frequency range, our results show viscous-fluid scaling of *G*″ at large *ω*, but experiments [22] and models [90] have shown an *ω*^3/4^ dependence at high frequencies. Now that we have performed a benchmark study which shows that a *non-Brownian* cross-linked actin network exhibits this behavior, it will be interesting to see if adding thermal bending modes [66, 10] reproduces the scaling relations observed previously, and how these relations are affected by hydrodynamic interactions. In concert with this, we will consider more bundled morphologies, where the flexibility of the filaments, and therefore their transverse bending fluctuations, may become more important. Similarly, we will test how explicit simulation of translational and rotational diffusion and (un)binding of individual CLs [66], as well as force dependence of CL kinetics, affect these results.

Other mechanical features of actin that will be interesting to explore include elastic twisting of actin filaments, which have been posited to contribute to elastic energy storage [52] and to the emergence of chirality in cross-linked gels [81]. Last, but not least, in vivo, the gels are very heterogeneous in the sense that there is a distribution of fiber lengths and turnover times, and a diversity of CL types co-existing with myosin motors [56, 91, 92]. How network heterogeneity and active molecular force generation affect the gel mechanics is another important question.

## Acknowledgements

Ondrej Maxian is supported by the National Science Foundation via GRFP/DGE-1342536 and Alex Mogilner is supported by National Science Foundation grant DMS1953430. This work was also supported by the NSF through Research Training Group in Modeling and Simulation under award RTG/DMS-1646339 and through the Division of Mathematical Sciences award DMS-2052515.

Code and input files for the simulations are available at https://github.com/stochasticHydroTools/SlenderBody. Supplemental movies are available at https://cims.nyu.edu/~om759/DynamicRheoVideos. All of our simulations were run on the NYU HPC Greene Supercomputer cluster.

## Footnotes

1 “Biologically relevant” timescales are from ~0.1 sec (the fastest relevant microscopic changes in the network structure, like unbinding of a CL [11]) to tens of seconds (turnover time, [27, 82]). The motility and division cycles for fast cells are on the scale of minutes (hundreds of seconds) [2]. Because of this, in vivo rheological experiments often focus on the range from 0.1 to 100 s [11] – we follow their example.

2 In experimental studies, what is measured is a *macroscopic* binding rate of the order 1 *μ*M^−1^ *s*^−1^, which cannot easily be converted to the effective microscopic rate we need. To find a suitable on rate, we make the observation from [86] that 10 – 50% of a 1 *μ*m long filament is decorated with *α*-actinin [86], which corresponds to about 1 – 10 CLs per filament of length 1 *μ*m; we find that *k*_on_ = 5/(*μ*m×s) is of the correct order of magnitude to give about 5 CL ends attached to each filament.

3 We obtained (19) by dropping the finite part integral in [7, Eq. (7.1)], which gives 1/2 instead of 3/2 in the denominator of [7, Eq. (8.13)]. Including intra-fiber hydrodynamics in the mobility changes the 0.5 in (19) back to 1.5. Substituting our parameters and recomputing, we get 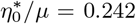 with intra-fiber hydrodynamics, which means the local drag approximation gives only 80% of the correct viscosity (see Section 3.7).

