## Supplemental figures for "Simulations of dynamically cross-linked actin networks: morphology, rheology, and hydrodynamic interactions"

### Supporting figures

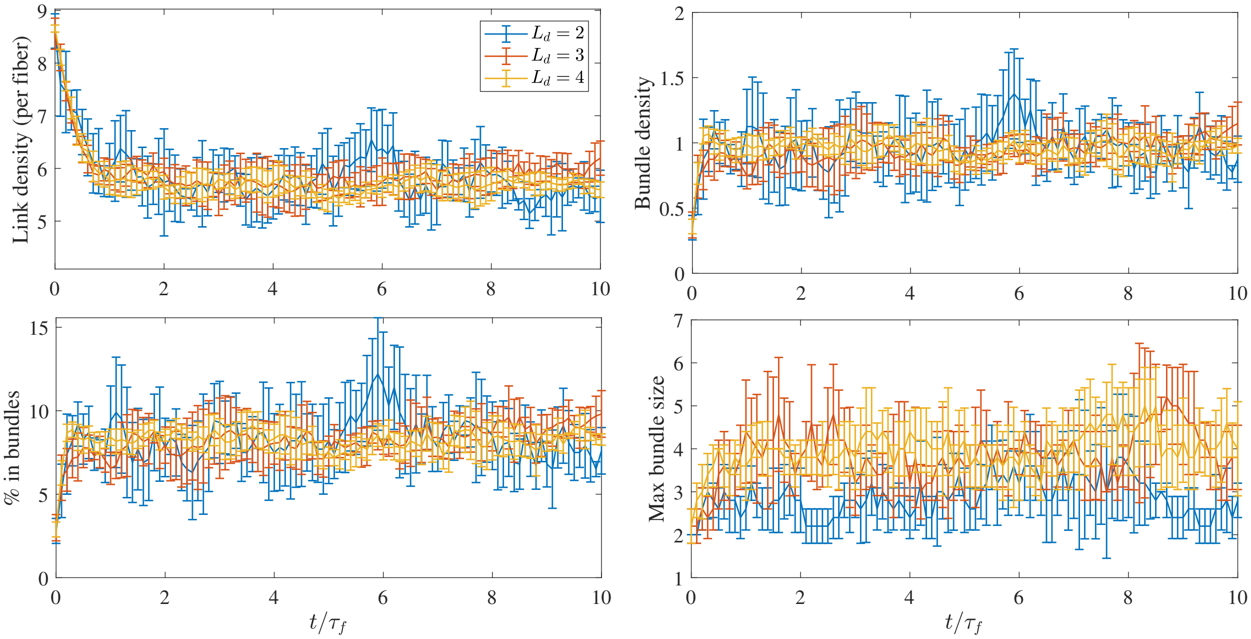


**Figure S1:** Number of links per fiber (top left), bundle density (top right, number of bundles / $L_d^3$), % of fibers in bundles (bottom left), and maximum bundle size (bottom right) for simulations from an initialized isotropic state run to the dynamic steady state for $\tau_f = 5$ seconds using three different system sizes: $L_d = 2$ (200 fibers, blue), $L_d = 3$ (675 fibers, red), and $L_d = 4$ (1600 fibers, yellow). Error bars are two standard errors over five trials.

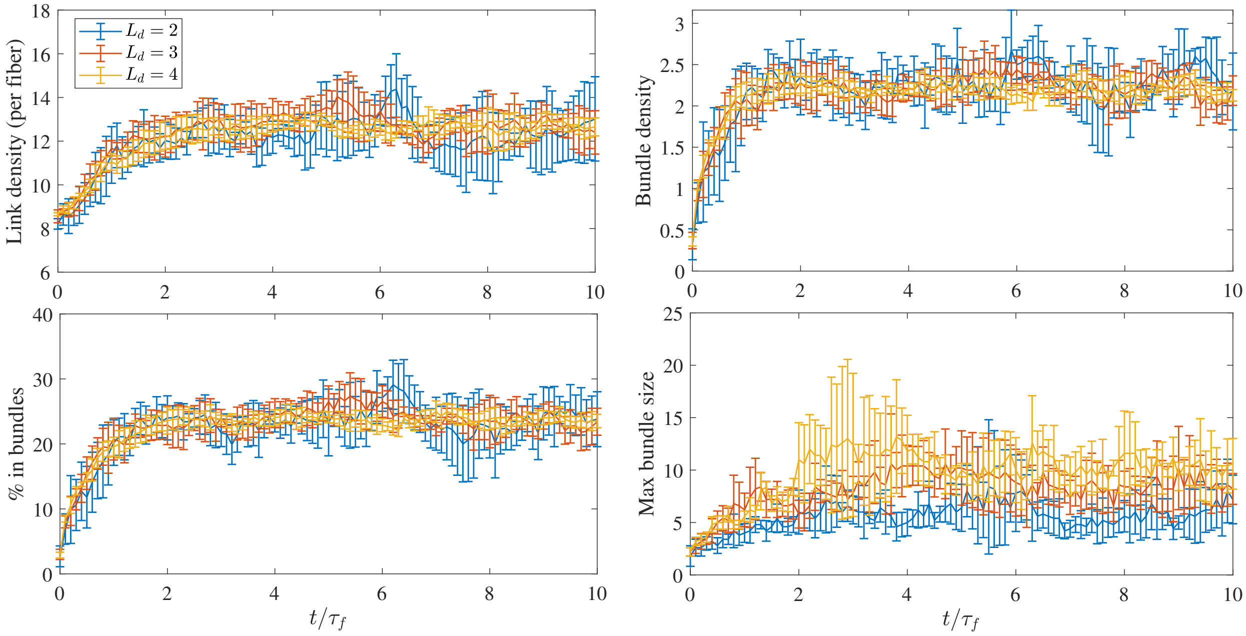


**Figure S2:** Number of links per fiber (top left), bundle density (top right, number of bundles / $L_d^3$), % of fibers in bundles (bottom left), and maximum bundle size (bottom right) for simulations from an initialized isotropic state run to the dynamic steady state for $\tau_f = 10$ seconds using three different system sizes: $L_d = 2$ (200 fibers, blue), $L_d = 3$ (675 fibers, red), and $L_d = 4$ (1600 fibers, yellow). Error bars are two standard errors over five trials.

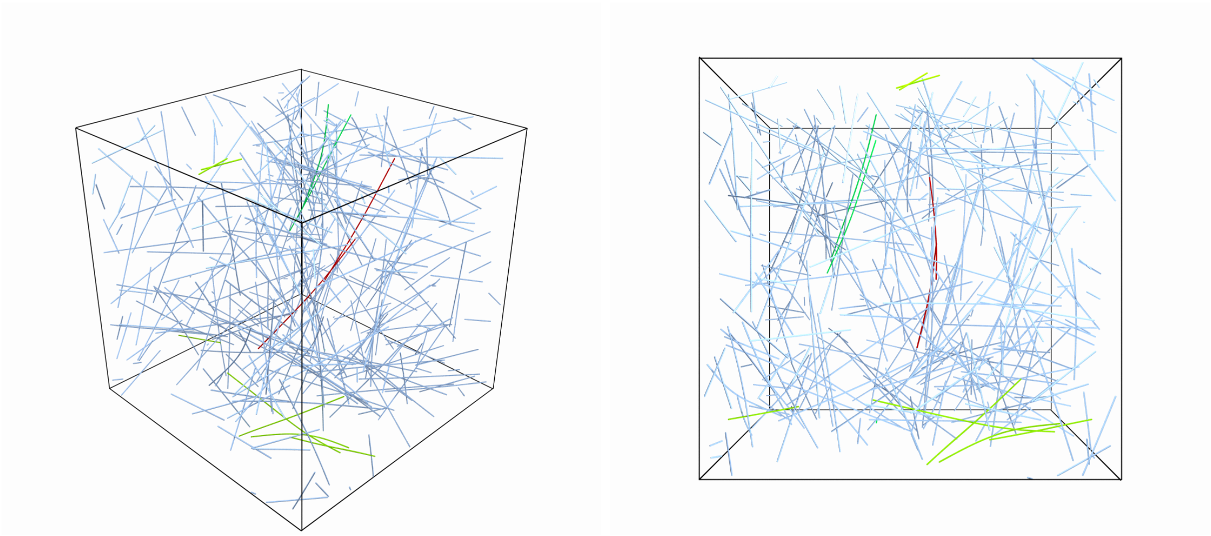

**Figure S3:** Side (left) and top (right) view of the homogeneous meshwork with $\kappa = 0.007$ pN$\cdot\mu$m$^2$, which is 1/10 of the stiffness of actin. Despite the smaller bending stiffness, the fibers still appear relatively straight, with the exception of some of the fibers in bundles.

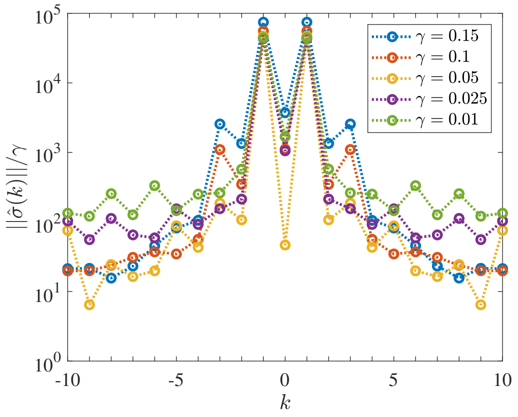


**Figure S4:** Spectrum of the stress for the B-In-M network. The coefficients $\hat{\sigma}(k)$ are defined in (18) in the main text. We consider various maximum strain values $\gamma = 0.01$ (green), 0.025 (purple), 0.05 (yellow), 0.1 (orange), and 0.15 (blue). We see the emergence of a $k = 3$ nonlinear harmonic for larger strains, suggesting a nonlinear response.

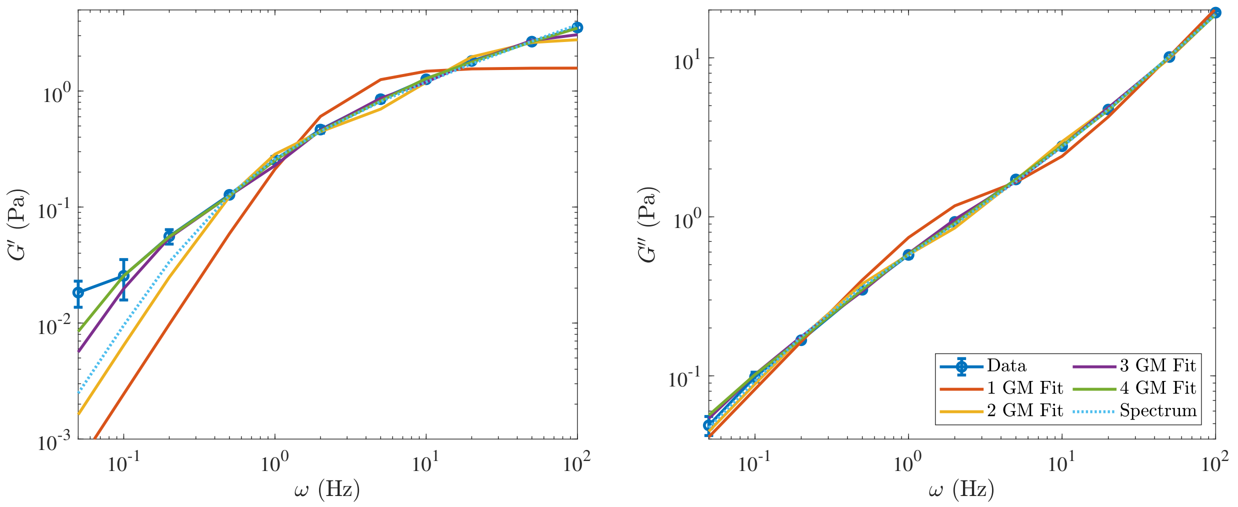


(a) B-In-M

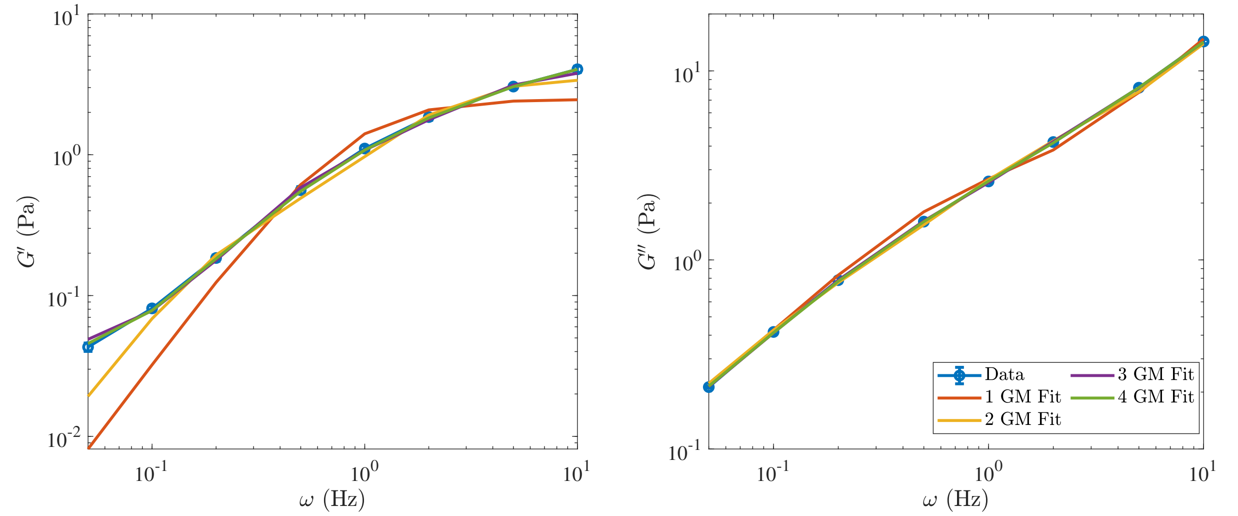


(b) B-In-M, $10\mu$

**Figure S5:** Fits to the elastic (left) and viscous (right) modulus data for (a) The B-In-M geometry with full hydrodynamics and (b) The B-In-M geometry with ten times larger viscosity and local drag mobility. We consider 1 (red), 2 (yellow), 3 (purple), and 4 (green) Maxwell modes in parallel with a viscous dashpot. The elastic modulus is only fit for $\omega \geq 0.1$ Hz. The B-In-M system with $\mu = 0.1$ Pa·s is best fit with three elements, while the higher viscosity system ($\mu = 1$ Pa·s) is best fit with only two elements, since in (b) the three and four element Maxwell models have a timescale larger than 100 seconds, indicating overfitting. For (a), the dashed line shows to a continuous spectrum of timescales, (25) in the main text, with $g_0 = 0.22$, $\alpha = 1.4$, $\tau_{\max} = 3.5$, and $\eta_0 = 0.16$, which is not a better fit than the 3 timescale generalized Maxwell model.
